# Non-CG DNA methylation and MeCP2 stabilize repeated tuning of long genes that distinguish closely related neuron types

**DOI:** 10.1101/2024.01.30.577861

**Authors:** J. Russell Moore, Mati T. Nemera, Rinaldo D. D’Souza, Nicole Hamagami, Adam W. Clemens, Diana C. Beard, Alaina Urman, Victoria Rodriguez Mendoza, Harrison W. Gabel

## Abstract

The extraordinary diversity of neuron types in the mammalian brain is delineated at the highest resolution by subtle gene expression differences that may require specialized molecular mechanisms to be maintained. Neurons uniquely express the longest genes in the genome and utilize neuron-enriched non-CG DNA methylation (mCA) together with the Rett syndrome protein, MeCP2, to control gene expression, but the function of these unique gene structures and machinery in regulating finely resolved neuron type-specific gene programs has not been explored. Here, we employ epigenomic and spatial transcriptomic analyses to discover a major role for mCA and MeCP2 in maintaining neuron type-specific gene programs at the finest scale of cellular resolution. We uncover differential susceptibility to MeCP2 loss in neuronal populations depending on global mCA levels and dissect methylation patterns and intragenic enhancer repression that drive overlapping and distinct gene regulation between neuron types. Strikingly, we show that mCA and MeCP2 regulate genes that are repeatedly tuned to differentiate neuron types at the highest cellular resolution, including spatially resolved, vision-dependent gene programs in the visual cortex. These repeatedly tuned genes display genomic characteristics, including long length, numerous intragenic enhancers, and enrichment for mCA, that predispose them to regulation by MeCP2. Thus, long gene regulation by the MeCP2 pathway maintains differential gene expression between closely-related neurons to facilitate the exceptional cellular diversity in the complex mammalian brain.

## Main

The development and function of the mammalian nervous system relies on a remarkably diverse array of cells with distinct functions within circuits. Single-cell transcriptomic studies have defined a hierarchy of thousands of cell types organized into neuronal versus non-neuronal cells, broad “classes” such as glutamatergic and GABAergic neurons, dozens of finer resolution “subclasses” made up of cells with related functions and developmental origins (e.g. somatostatin (SST)- and parvalbumin (PV)-positive inhibitory interneurons), and hundreds of closely related cell “types” with putatively distinct functional roles in circuits (e.g. subsets of PV neurons)^1^. Notably, at the highest level of resolution, many neuron types emerge in postnatal development, and the development of distinct genetic programs can be influenced by environmental stimuli^2,3^. These types are often distinguished not by unitarily expressed marker genes, but rather through subtly tuned differences in mRNA expression across numerous genes. The existence of this extreme diversity derived from small differences in many genes in the late stages of development raises questions of how the mammalian nervous system can stably maintain transcriptomic differences across closely related types throughout the lifetime of the organism, and whether specialized gene regulatory mechanisms are necessary to facilitate this process. Furthermore, while numerous types can be defined by measuring single-cell gene expression, the functional importance of maintaining these high-resolution gene expression differences is debated, and it is not clear if disruption of these subtle differences contributes to dysfunction in disease states.

DNA methylation in neurons displays unique characteristics that suggest that this epigenetic mark supports transcriptomic diversity across cell types^4^. While all mammalian cells utilize cytosine methylation at CG dinucleotides (mCG) to control transcription, neurons contain high levels of non-CG methylation not present in other cell types. Non-CG methylation primarily occurs at CA dinucleotides (mCA) and accumulates in the early postnatal period until this unique mark accounts for approximately half the total methylation sites in neurons^5^. High levels of mCA are a hallmark of vertebrate neurons, suggesting that this epigenetic mark has a unique role in the development of complex nervous systems^6^. Global levels of mCA vary across different brain regions and cell types, and patterns of mCA across genes display strikingly cell-type-specific patterning^7–9^, suggesting an important role in regulating cell-type-specific transcription. Despite these observations, there have been limited studies to date demonstrating the mechanism and impact of mCA-mediated gene regulation at the level of individual subclasses or high-resolution cell types.

The methyl-CpG binding protein 2 (MeCP2) has been identified as a critical reader of mCA, binding to this mark and canonical mCG to repress transcription. A major target of MeCP2 regulation is long genes (e.g. genes longer than 100 kb) that are embedded in regions of high mCA^10–16^. MeCP2 has been shown to control these genes, at least in part, by recruiting the NCoR co-repressor complex^17^ and repressing the activity of enhancers^12,18^, particularly intragenic enhancers found within introns^12,19^. Loss or overexpression of MeCP2 leads to subtle but widespread effects across many genes, suggesting that this protein functions with DNA methylation to tune transcription in the brain rather than turn genes off and on^20,21^. These studies have revealed mechanisms by which MeCP2 can work with mCA to regulate genes, but how MeCP2 reads distinct methylation patterns in each cell type and whether it contributes to differential gene expression across closely related cell types have not been defined.

As the major reader of cell-type-specific mCA, loss of MeCP2 might be expected to impact different genes across distinct brain regions and cell types, and indeed some studies of MeCP2 mutants performed across different brain regions and broad cell classes (e.g. inhibitory and excitatory neurons) have focused on unique MeCP2 gene targets in each dataset^22–24^. Paradoxically, however, other MeCP2 mutant studies have detected sets of dysregulated genes that overlap across brain regions and cell populations^10,13,25^. These core MeCP2-repressed genes (those upregulated in *Mecp2*-null mutants) are very long genes, with annotations of key functions in neurons^13,26^, suggesting that their regulation is critical to nervous system function. How these genes could be consistently impacted across brain regions and neuron classes despite the cell-type-specific nature of mCA has not been resolved, and the functional significance of this pathway preferentially targeting long genes is not known. Together, these apparently discordant results make it unclear if the MeCP2 pathway regulates a core set of genes across all cell types, or if it defines cell-type-specific gene programs.

Deciphering the overarching biological impact of the MeCP2 pathway across cell types is critical, as disruption of this pathway is implicated in multiple neurodevelopmental disorders. Loss of mCA through mutation of the de novo methyltransferase DNMT3A causes severe neurological phenotypes in mice, and disruption of DNMT3A itself, or molecular pathways that target DNMT3A to methylate the neuronal genome, leads to Tatton-Brown-Rahman syndrome and Sotos syndrome^27–30^. Furthermore, loss or overexpression of MeCP2 causes Rett syndrome (RTT) or MeCP2 duplication syndrome (MDS), respectively^31–33^. Rett syndrome is typified by postnatal onset, and the accumulation of mCA and MeCP2 to high levels in the brain during a dynamic period of postnatal circuit refinement and critical period closure suggests an important role for this pathway in defining mature circuits in the brain. However, the mechanisms by which disruption of mCA and MeCP2 drive dysfunction in disease, and whether this pathology arises from broad dysregulation of a core set of genes across many cell types, or through distinct dysregulation in each cell type, remains to be determined.

Here we address these outstanding questions by employing cell-type-specific epigenomic profiling and spatial transcriptomics to dissect gene regulation by mCA and MeCP2 across neuron populations. We demonstrate that high mCA cell populations are more susceptible to gene dysregulation upon loss of MeCP2 than those with low mCA cells and show that megabase-scale patterns of mCA enrichment drive overlapping enhancer and gene regulation by MeCP2 across neuron populations, while cell-specific gene body depletion of mCA is linked to differential MeCP2-mediated repression between populations. Strikingly, we discover that a major function of these MeCP2 regulatory mechanisms is to stabilize the differential expression of genes that distinguish neuron types at the highest resolution. We show that long, type-specific genes with essential roles in neurons are often “repeatedly tuned” between different groups of highly-related neuron types in distinct subclasses (e.g., types found within distinct inhibitory and excitatory neuron subclasses), and that these repeatedly tuned genes display DNA methylation patterns and gene regulatory structures that predispose them to regulation by mCA and MeCP2 across cell types. We further show that this tuning function for MeCP2 is required to maintain spatially resolved gene expression patterns that emerge in sublayers of the visual cortex during postnatal development. Together, our findings suggest that mCA and MeCP2 in vertebrate neurons is a key pathway to maintain the extreme cellular transcriptomic diversity required in complex neural circuits, and implicate disruption of fine-scale gene expression tuning across closely related neuron types in the pathology of neurodevelopmental disorders.

## Results

### Global mCA levels determine the overall impact of MeCP2 in distinct neuronal populations

While major differences in levels and patterns of mCA exist between brain regions and cell types, the quantitative impact of these differences on MeCP2 gene regulation has not been systematically explored. We therefore started our investigation of neuron type-specific MeCP2 function by examining effects of MeCP2 loss across subclasses of neurons with distinct global levels of mCA. We hypothesized that upon MeCP2 loss, cell populations with high mCA levels should display more transcriptional dysregulation than those with lower levels of this methyl mark. We utilized the Isolation of Nuclei Tagged in specific Cell Types methodology (INTACT)^9,34^ to profile four neuron subclasses in the cerebral cortex with diverse physiologic functions and varying global mCA levels^8^ (Fig. 1a): somatostatin-positive (SST) and fast-spiking parvalbumin (PV) interneurons, which are enriched for mCA and implicated in Rett-like pathology in mice ^35,36^; as well as layer 4 (L4) and layer 5 (L5) excitatory neurons which show low and intermediate levels of mCA, respectively^8^ (Extended Data Fig. 1a). We collected nuclei from MeCP2 KO and WT littermate pairs for RNA-sequencing. Similar numbers of each subclass were isolated between MeCP2 KO and WT mice (Fig. 1b and Extended Data Fig. 1b), and marker gene expression validated specificity (Fig. 1c and Extended Data Fig. 1c). To analyze the correlation between mCA and gene expression effects, we compiled and validated methylomes for each subclass by merging single-nucleus methylation data from the 8-week mouse cortex^8^. Each of these aggregate methylomes displayed the known anticorrelation between mCA levels in genes and subclass-specific gene expression, validating the concordance between our datasets (Extended Data Fig. 1d,e).

**Fig. 1.**
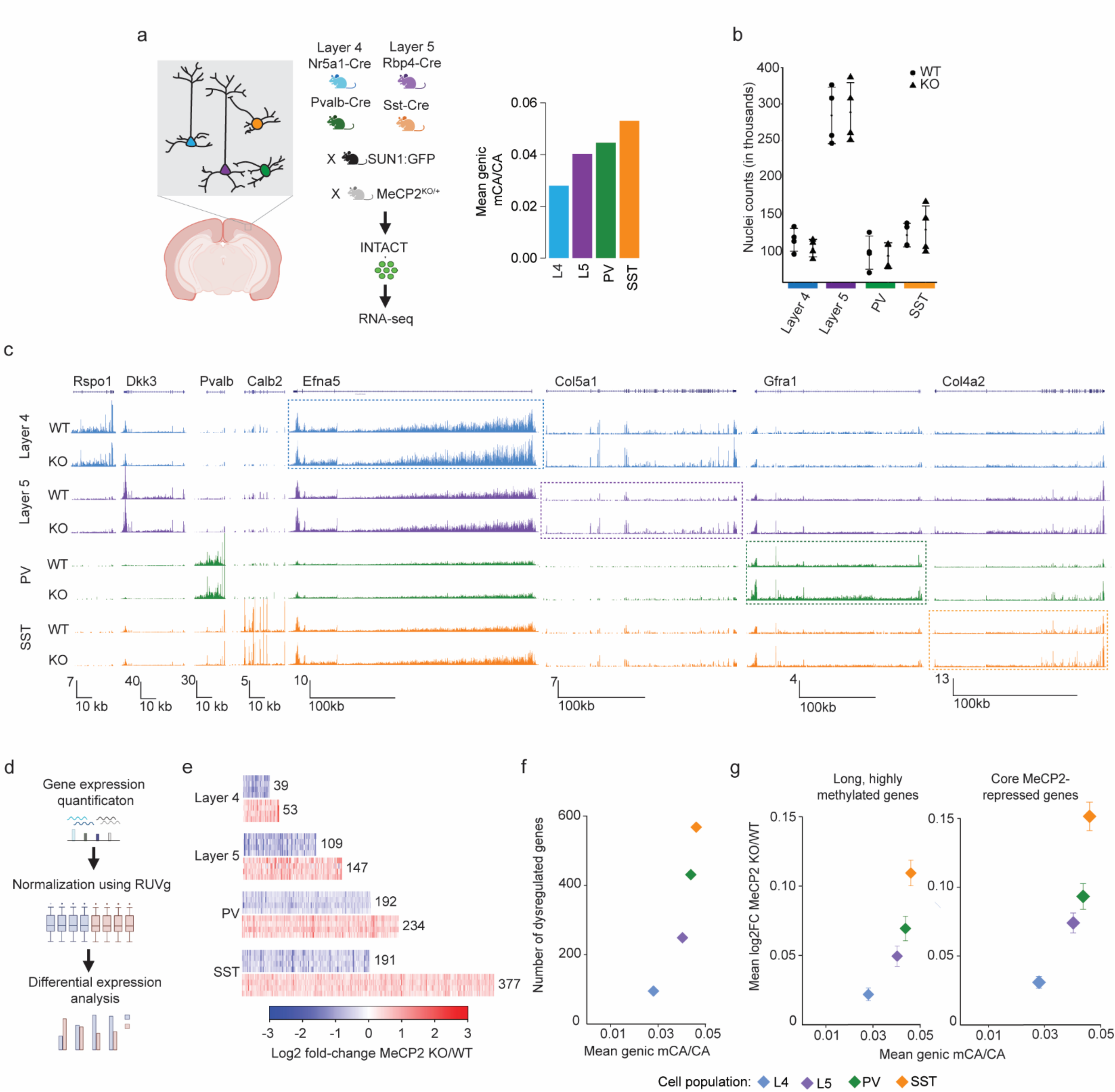
Neuron subclass-specific analysis of global mCA levels and gene dysregulation in the MeCP2 KO. **a,** Four neuron subclasses from the cerebral cortex with varying levels of mCA were selected for gene expression analysis in MeCP2 KO using the INTACT nuclear isolation system. **b,** Number of nuclei isolated by INTACT for KO and WT animals for each population profiled. No significant differences in numbers of nuclei isolated were detected. **c,** Genome browser view of RNA-seq data for MeCP2 WT and KO in each population. Marker genes *Rspo1, Dkk3, Pvalb,* and *Calb2* for each subclass profiled (L4, L5, PV, and SST, respectively) shown to the left. Example MeCP2-repressed gene for each subclass is highlighted with a box (*Efna5* – L4, *Col5a1* – L5, *Gfra1* – PV, *Col4a2* – SST). **d,** Overview of analysis workflow of differential gene expression analysis for INTACT-RNA-seq. **e.** Heatmap of per replicate fold-changes MeCP2 KO/WT of significantly down-regulated genes (blue) and up-regulated (red) genes identified in each subclass with total number of genes shown. **f,** Scatter plot of the number of significantly dysregulated genes identified in each subclass plotted vs average mCA level in that subclass. **g**, Scatter plot of mean fold-change of long (greater than 100 kb), highly methylated (top decile of mCA) genes (left) and core MeCP2-repressed genes (right) vs. average mCA level for all genes in each subclass. n=4 biological replicates for PV, SST, L4, and L5 RNA-seq.

To interrogate the association between mCA levels and the magnitude of MeCP2 gene regulation in neuron subclasses, we performed RUVg normalized^37^ differential expression analysis between MeCP2 KO and WT cells in each subclass (Fig. 1d). This analysis revealed a clear correlation between the number of dysregulated genes and mCA levels in each subclass (Fig. 1e,f). Gene expression studies have revealed subtle sub-significance threshold effects in which MeCP2 generally represses longer genes containing high levels of mCA^10,12,15,23^, and we find that mCA levels in each subclass correlate with the effect size for the longest (>100 kb), most highly methylated genes mCA (top 10% highest methylated) in that subclass (Fig. 1g). We also observed greater overall spread in gene fold-changes in higher mCA subclasses genome-wide (Extended Data Fig. 1f). We further assessed the effect on a set of genes identified as MeCP2-repressed across studies of multiple brain regions^13^. These “core MeCP2-repressed genes” are long and highly methylated in all data sets examined to date^13^ (Extended Data Fig. 1g), and approximately 80% of these genes are upregulated in any given MeCP2 KO dataset^12,13^. We find similar correlation between cellular mCA levels and the magnitude of dysregulation for this gene set (Fig. 1g). Finally, analysis of bulk RNA-sequencing from brain regions revealed similar correlations between effect sizes and global levels of mCA (Extended Data Fig. 2a-c). Together, these data indicate that the level of mCA in distinct populations of neurons correlates with the magnitude of dysregulation when MeCP2 is lost. Thus, cells with the highest levels of mCA are likely most affected by loss of mCA or MeCP2 in disease, and may have the largest influence on pathology of Rett syndrome and other disorders involving the MeCP2 pathway.

### Regional and gene-specific mCA patterning drive overlapping and distinct regulation by MeCP2

The nature of the gene programs targeted by MeCP2 has been debated, with some previous studies of gene expression upon MeCP2 loss emphasizing nonoverlapping dysregulated gene sets across cell types and brain regions^23,24^ and others highlighting overlapping gene targets, such as core MeCP2-repressed genes^12,13,38^. To address these apparently incongruous findings, we assessed overlapping and distinct MeCP2 gene regulation between neuronal subclasses and explored methylation patterns that could drive these effects. This analysis revealed significant shared gene dysregulation across all comparisons (Fig. 2a-b and Extended Data Fig. 3a). Additionally, we observed high overlap between the MeCP2-repressed gene list in each subclass and the core MeCP2-repressed genes previously identified across non-cortical brain regions (Fig. 2a-b). In the context of this extensive overlap, we did detect genes that are uniquely dysregulated in each subclass, demonstrating that MeCP2 can mediate distinct as well as overlapping transcriptomic regulation (Fig. 2a). These findings indicate that across subclasses, a major subset of genes is predisposed to repression by MeCP2, but that in each subclass, distinct gene regulation can also occur.

**Fig. 2.**
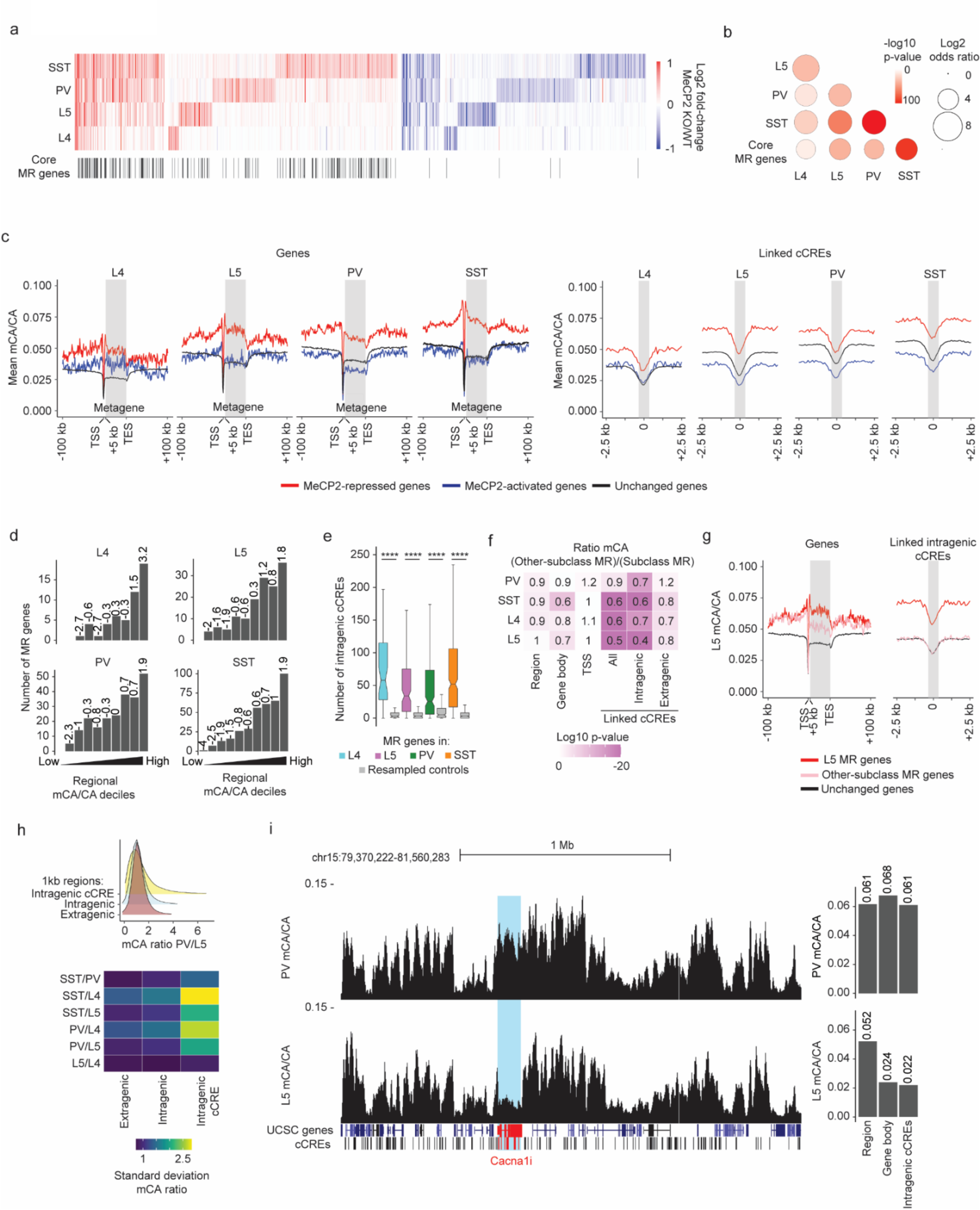
Overlapping and distinct regulation of genes across neuronal populations by MeCP2 is associated with regional and gene-specific mCA patterns. **a,** Heatmap of fold-changes for overlapping and unique significantly dysregulated genes in each neuronal subclass. Core MeCP2-repressed (MR) genes marked in black below. **b,** Significance of overlap of MeCP2-repressed genes in each subclass and core MeCP2-repressed genes from multiple datasets. P-values are calculated by two-sided Fisher’s exact test. **c,** Left: aggregate mCA/CA levels for MeCP2-regulated genes identified in L4, L5, PV, and SST neurons. Mean mCA/CA is reported for 1 kb bins. “Metagene’’ refers to 50 equally sized bins within gene bodies. Right: aggregate mCA/CA levels centered at cCREs linked to MeCP2-regulated genes in L4, L5, PV, and SST neurons. Mean mCA/CA is reported for 100 bp bins. Gray rectangle = 700 bp, ~ median length of all cCREs. **d,** Number of MeCP2-repressed genes found in each decile of genes sorted by regional mCA level. Regional mCA for each gene is calculated as mCA/CA for the region 10 kb to 210 kb upstream of the TSS and the region 200 kb downstream of its TES, removing the signal from genes overlapping these regions. Enrichment (log2 odds ratio) of MeCP2-repressed genes in each decile shown above each count for each decile. **e,** Number of intragenic cCREs inside MR genes identified in L4, L5, SST, and PV neurons and expression-resampled controls. The center line is the median. Each box encloses the first and third quartile of the data. The whiskers extend to the most extreme values, excluding outliers which are outside 1.5 times the interquartile range. ****p < 0.0001 two-sided Wilcoxon rank-sum test. **f,** Heatmap of mCA/CA enrichment in regions, gene bodies, and linked cCREs of other-subclass MR genes over those of subclass MR genes, colored by the log10 two-sided Wilcoxon rank-sum p-value. Numbers in the tiles represent the ratio of median mCA/CA of elements associated with subclass MR genes to the median mCA/CA of elements associated with other-subclass MR genes. **g,** Aggregate mCA/CA levels at gene bodies (left) and linked cCREs (right) of L5 MR genes, other-subclass MR genes, and unchanged genes. **h**, Top: density plot of mCA/CA ratio between PV and L5 neurons in 1 kb extragenic regions, intragenic regions, and regions centered at intragenic cCREs. Bottom: heatmap of standard deviation mCA/CA ratios between pairs of subclasses among L4, L5, PV, and SST neurons. **i**, Left: genome browser view of PV (top) and L5 (bottom) mCA/CA at *Cacna1i*, a gene repressed by MeCP2 in PV neurons but not in L5 neurons. Right: regional, gene body, and intragenic linked cCRE mCA levels of *Cacna1i* in PV (top) and L5 (bottom) neurons. Numbers on top of bars are the mCA/CA levels of each group. DNA methylation data were compiled from previously published single-cell methylomic analysis^7^, *see methods*. RNA-seq data are from INTACT analysis described in **Figure 1**, n=4 biological replicates per genotype per cell type.

To understand how DNA methylation drives significant overlapping MeCP2 regulation across subclasses while also impacting distinct genes in each subclass, we performed integrated analysis of gene expression effects and DNA methylation patterns across the genome. Tissue-level studies have shown that many MeCP2-repressed genes, putative direct targets of repression by the MeCP2-NCoR complex^39,40^, are embedded in megabase-scale regions of the genome that accumulate high mCA. This high regional mCA “set-point” leads to enrichment of mCA within gene bodies and at enhancers associated with these genes. The enrichment of mCA at enhancers, particularly intragenic enhancers found inside the gene, is associated with repression by MeCP2^12,27^. Notably, however, genes can opt-out from high methylation if they are highly expressed during the early postnatal period, as deposition of gene body histone modifications associated with transcription can block recruitment of DNMT3A and lead to lower accumulation of mCA within the gene^27,41^. While this two-step, regional set-point and gene-specific depletion patterning mechanism for mCA has been described in brain tissue, the contributions of these steps to overlapping or distinct gene regulation by MeCP2 in individual neuron subclasses has not been assessed. We therefore evaluated methylation in and around MeCP2-repressed genes and at putative enhancers, or candidate cis-regulatory elements (cCREs), previously linked to these genes by single-cell ATAC-seq of the adult mouse brain^42^.

In each subclass, we find that MeCP2-repressed genes are embedded in regions of high mCA resulting in high gene body and cCRE mCA levels (Fig. 2c,d and Extended Data Fig. 3b,e). Furthermore, MeCP2-repressed genes in each subclass are long and contain many intragenic enhancers (Fig. 2e and Extended Data Fig. 3f), which show the highest mCA enrichment of all elements associated with MeCP2-repressed genes (Extended Data Fig. 3b). Subtle, but significant mCG enrichment is also present at linked cCREs (Extended Data Fig. 3c,d). In contrast to MeCP2-repressed genes, genes that show a relative reduction in expression upon loss of MeCP2 (“MeCP2-activated” genes) tend to be depleted for methylation at their linked cCREs relative to unchanged genes (Fig. 2c and Extended Data Fig. 3b-d), suggesting that these genes escape repression by MeCP2. These observations support a model in which regional mCA levels in each cell population influence enhancer methylation, and mCA levels at these enhancers, particularly intragenic enhancers, in turn drive the extent of repression by MeCP2.

If high regional mCA predisposes intragenic enhancers and their target genes to MeCP2-mediated repression in each cell type, what might prevent genes in high-mCA regions from being repressed by MeCP2 in some specific subclasses? To address this, we assessed how subclass-specific methylation profiles vary across regions, gene bodies, and enhancers associated with genes that are differentially impacted by MeCP2 between subclasses. Comparison of genes that are significantly repressed in a given cell subclass (“subclass MeCP2-repressed” genes) to genes that are not significantly repressed by MeCP2 in that subclass but are MeCP2-repressed in other subclasses (“other-subclass MeCP2-repressed” genes), revealed that gene body and cCRE methylation was depleted in other-subclass MeCP2-repressed genes relative to subclass MeCP2-repressed genes (Fig. 2f,g). Notably, the strongest signal for differential methylation occurs at intragenic cCREs linked to MeCP2-repressed genes (Fig. 2f), implicating mCA at these sites as a major driver of subclass-specific regulation by MeCP2. In contrast, subclass MeCP2-repressed genes and other-subclass MeCP2-repressed genes show similar regional mCA levels (Fig. 2f,g). Examination of methylation associated with core MeCP2-repressed genes that escape repression in each neuron subclass revealed similar mCA patterns (Extended Data Fig. 3g-i). Together, these results support the model in which regional mCA predisposes genes to MeCP2-mediated repression, but selective protection from methylation in a given subclass can exclude these genes from repression to solidify subclass-specific gene expression programs.

The observation that high regional mCA for MeCP2-repressed genes is invariant across subclasses while gene body and cCRE mCA at these genes can be selectively depleted suggests that these two patterns of methylation may fundamentally differ in their subclass-to-subclass variation genome-wide. Indeed, comparison of 1 kb windows across the genome revealed that intragenic regions show the greatest degree of variation in mCA between subclasses compared to extragenic regions (Fig. 2h and Extended Data Fig. 3j). For example, *Cacna1i*, which is repressed by MeCP2 in PV cells but not in L5 excitatory cells, is embedded in a region with high mCA in both PV and L5 cells, but the transcribed region and associated cCREs of *Cacna1i* are lowly methylated in L5 cells relative to PV cells (Fig. 2i). Thus, while regional methylation varies widely between different large-scale regions in the genome, the relative enrichment of mCA for a given region is similar between subclasses. This cell-type-invariant regional enrichment of mCA explains how MeCP2-repressed genes are found in regions of high mCA in all cell types and therefore predisposed to MeCP2 regulation.

### MeCP2 represses enhancers from non-cognate cell types to tune gene expression

To investigate the mechanism by MeCP2 and DNA methylation regulate gene expression in an individual subclass, we next carried out subclass-specific epigenomic profiling. For this analysis, we focused on PV interneurons, and utilized the INTACT system to assess differential enhancer activation^43^ by ChIP-seq analysis of histone H3 lysine 27 acetylation (H3K72ac). To focus our analysis on the direct effects of MeCP2, we conducted combined analysis on INTACT-isolated PV nuclei from MeCP2 KO and MeCP2 overexpression (OE) mice (Fig. 3a) and searched for reciprocal effects in these mutants. H3K27ac ChIP-seq signal in isolated nuclei displayed strong differential signal at genes with PV-specific expression compared to total cortical nuclei, verifying specificity for PV neurons (Fig. 3b-c).

**Fig. 3.**
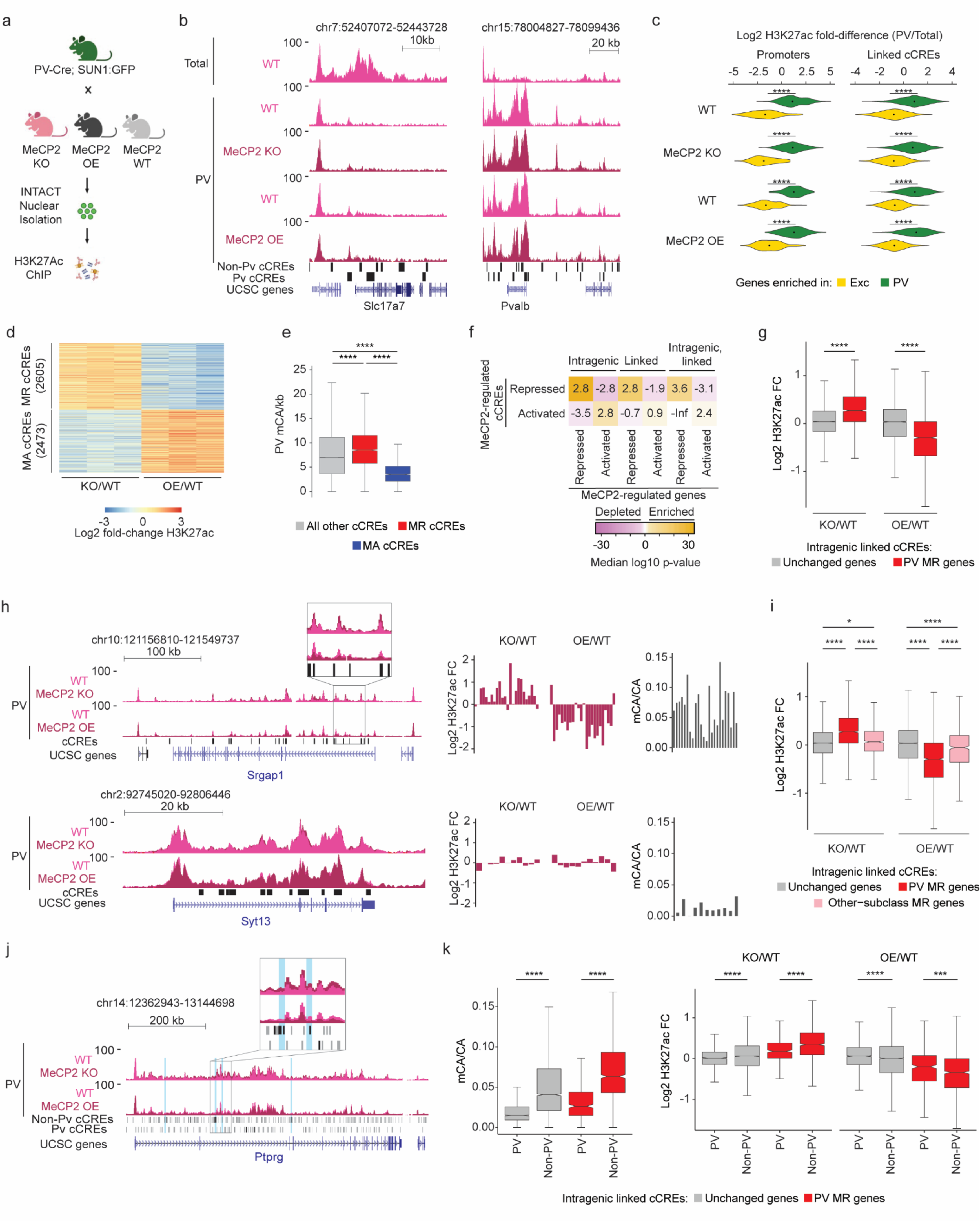
Dysregulation of cell-type-specific high mCA enhancers in MeCP2 knockout parvalbumin positive interneurons. **a,** Schematic of INTACT isolation and H3K27ac ChIP-seq profiling of PV MeCP2 wild-type (WT), MeCP2 knockout (KO), and MeCP2 overexpression (OE) neurons. **b,** Genome browser views of H3K27ac ChIP-seq signal in cortical (total) and PV nuclei at *Slc7a7*, a gene expressed in excitatory neurons, and *Pvalb*, a marker gene for PV neurons. Location of cCREs and genes shown at bottom. **c**, Log2 H3K27ac fold difference between PV and total nuclei for the same genetic background (MeCP2 WT, MeCP2 KO, or MeCP2 OE) at promoters and linked cCREs of the top 100 genes enriched for expression in PV neurons (log2 gene expression difference (PV/excitatory) > 0, PV RPKM >= 10, FDR-adjusted p-value <= 0.01, ordered by log2 fold difference) and the top 100 genes enriched in excitatory (Exc) neurons (log2 gene expression fold difference (excitatory/PV) > 0, Exc RPKM >= 10, FDR-adjusted p-value <= 0.01, ordered by log2 fold difference). The genes were identified from previous differential gene expression analysis of PV and excitatory genes^52^. ****p < 0.0001 two-sided Wilcoxon rank-sum test. Violin plots were made after excluding outliers which are outside 1.5 times the interquartile range of the data. **d,** H3K27ac fold-changes for cCREs identified as significantly dysregulated for H3K27ac ChIP-seq signal in PV neurons isolated from MeCP2 KO and MeCP2 OE mice (FDR-adjusted p-value ≤ 0.1). MR = MeCP2-repressed; MA= MeCP2-activated. **e,** Boxplot of PV mCA/kb in MeCP2-repressed (MR), MeCP2-activated (MA) and all other cCREs. ****p < 0.0001 two-sided Wilcoxon rank-sum test. **f,** Quantification of enrichment of MeCP2-regulated cCREs to be located inside (intragenic) or linked to MeCP2-regulated genes by co-correlation analysis^42^ or both. Median significance (log10 p-value from two-sided Fisher’s exact test, color) and enrichment (log2 odds ratio, number) are shown for cCREs associated with MeCP2-regulated genes compared to cCREs associated with expression-resampled genes. **g,** Log2 H3K27ac fold-change for cCREs located inside of, and linked to, PV MR genes or unchanged genes in PV MeCP2 KO and MeCP2 OE. ****p < 0.0001 Two-sided Wilcoxon rank-sum test. **h,** Left: overlaid PV MeCP2 WT, MePC2 KO, and MeCP2 OE H3K27ac ChIP-seq tracks in the PV MeCP2-repressed gene *Srgap1* (top) and an other-cell-type MeCP2-repressed gene *Syt13* (bottom). The inset shows cCRE-containing regions with changes in H3K27ac upon MeCP2 perturbation. Right: log2 H3K27ac fold-change PV MeCP2 KO and MeCP2 OE and PV mCA/CA of cCREs inside and linked to *Srgap1 or Syt13*. The expression of *Syt13* is not affected in PV neurons and it does not show enrichment of mCA or alterations in histone acetylation at its associated cCREs. **i,** Log2 H3K27ac fold-change of cCREs inside and linked to PV MR genes, other-cell-type MR genes, or unchanged genes in PV MeCP2 KO and MeCP2 OE. *p < 0.05, ****p < 0.0001 Two-sided Wilcoxon rank-sum test. **j,** Genome browser view of H3K27ac ChIP-seq at the *Ptprg* gene in PV wild-type, MeCP2 KO, and MeCP2 OE. Gray bars are all cCREs while black bars are intragenic, cCREs linked to the gene. Inset shows region containing non-PV cCREs with changes in H3K27ac ChIP-seq signal in MeCP2 KO and MeCP2 OE relative to wild-type. Blue highlights are examples of non-PV cCREs called as significantly dysregulated upon MeCP2 perturbation. **k,** PV mCA/CA (left) and log2 H3K27ac fold-change in PV MeCP2 KO and MeCP2 OE (right) of PV and non-PV cCREs located inside of, and linked to, unchanged genes or PV MR genes. ***p < 0.001, ****p < 0.0001 two-sided Wilcoxon rank-sum test. n=3 biological replicates for PV WT, MeCP2 KO, and MeCP2 OE H3K27ac ChIP-seq. n=2 biological replicates for Total WT, MeCP2 KO, and MeCP2 OE H3K27ac ChIP-seq. For all boxplots, the center line is the median. Each box encloses the first and third quartile of the data. The whiskers extend to the most extreme values, excluding outliers which are outside 1.5 times the interquartile range.

Differential analysis of H3K27ac ChIP-seq signal at cCREs in MeCP2 KO and MeCP2 OE PV nuclei using edgeR^44^ (Fig. 3d), identified 5078 MeCP2-regulated cCREs. We classified cCREs significantly increased H3K27ac signal in the MeCP2 KO and decreased H3K27ac signal in the MeCP2 OE as MeCP2-repressed cCREs, and those significantly affected in the opposite direction as MeCP2-activated cCREs. Consistent with a role for MeCP2 in enhancer repression in PV neurons, we found that MeCP2-repressed cCREs are enriched for mCA, mCG, and MeCP2 binding, and are located within genes more than MeCP2-activated cCREs and cCREs not significantly affected (Fig. 3e and Extended Data Fig. 4a-c). We find that these altered enhancers are located within and are functionally linked to MeCP2-repressed genes (Fig. 3f). Furthermore, analysis of all cCREs linked to MeCP2-repressed genes revealed upregulation of H32K7ac signal in the MeCP2 KO and downregulation of H3K27ac signal in the MeCP2 OE at these sites (Fig. 3g). These results demonstrate enhancer regulation by MeCP2 in PV neurons is directly linked to control of gene expression within these cells.

We next interrogated the role of differential enhancer regulation in subclass-specific gene repression by MeCP2. Consistent with our differential DNA methylation analysis above (e.g., Fig. 2f) we find that intragenic cCREs linked to PV MeCP2-repressed genes are enriched for MeCP2 binding and display robust changes in H3K27ac in MeCP2 mutants, while cCREs linked to other-subclass MeCP2-repressed genes not affected in PV cells display limited MeCP2 binding and H3K27ac fold-changes (Fig. 3h-i, Extended Data Fig. 4d,e). These data suggest that MeCP2 regulates genes in a subclass-specific manner by reading out enhancer methylation patterns in each subclass and differentially repressing enhancer activation.

Cell-type-specific enhancer activation is key to coordinating cell-type-specific gene expression programs, and DNA methylation may represent a critical mechanism to maintain cell-type specificity of enhancer activation^45,46^. We therefore investigated if loss of MeCP2 impacts repression of enhancers that are not normally highly active in PV neurons. Indeed, visualization of PV H3K27ac signal at genes and enhancers revealed examples of acetylation changes in regions with low baseline H3K27ac in wild-type samples that might represent de-repression of cCREs that are robustly active in other cell types but not normally active in PV neurons (Fig. 3j). Splitting cCREs into those active in PV neurons (PV-cCREs) in single cell ATAC-seq analysis^42^ compared to cCREs not detected as active in these cells (non-PV cCREs) and assessing their characteristics, we found that non-PV cCREs linked to PV MeCP2-repressed genes display greater methylation, MeCP2 binding, and H3K27Ac fold-changes than PV cCREs linked to MeCP2-repressed genes (Fig. 3k and Extended Data Fig. 4f). HOMER analysis^47^ detected transcription factor motif enrichment in non-PV MeCP2 repressed cCREs corresponding to transcription factors that are expressed in PV neurons and play important roles in neuronal cell type diversity (e.g., the ROR family)^7,48^ (Extended Data Fig. 4g and Table S1). This suggests that loss of MeCP2 in PV neurons creates a permissive environment that allows these transcription factors to spuriously activate enhancers normally kept off in PV neurons by DNA methylation. Together, these findings indicate that a major target of repression by MeCP2 in PV neurons is enhancers that are most active in other subclasses, suggesting that downregulation of these enhancers by MeCP2 helps to define gene expression patterns within specific neuronal populations.

### The MeCP2 pathway regulates genes that are repeatedly tuned across closely related neuron types

Our analyses shed light on the mechanism by which MeCP2 and mCA regulate transcriptional programs in subclasses of neurons, but the overarching biological function of this unique regulatory pathway has remained elusive. We therefore examined the functional annotations and expression patterns of MeCP2-regulated genes we identified in each subclass, searching for unifying characteristics. Gene ontology analysis of MeCP2-repressed genes identified key functional annotations of genes that were disrupted across subclasses, including ion channels, cell-adhesion molecules, and extracellular matrix proteins (Fig. 4a and Extended Data Fig. 5a). Notably, enriched functional categories in each subclass showed substantial overlap, suggesting that genes that precisely modulate the connectivity and physiology of neuronal types are commonly fine-tuned by mCA and MeCP2. These genes are important for the physiology in the neuronal subclasses profiled. For instance, in PV interneurons, we find that sets of ion channel genes, including *Kcnh5* and *Kcns1*, known to contribute to the firing properties of these neurons^49,50^, are repressed by MeCP2. We further noted that genes we detect as MeCP2-regulated in each subclass are important for defining the functional identities of PV and SST neurons, including higher resolution neuronal types within these subclasses. These include the metabotropic glutamate receptor subtype 1 gene *Grm1*; *Lypd6*, a modulator of nicotinic receptor function; and *Cdh13*, which plays a critical role in maintaining the excitation/inhibition balance within brain networks^51–54^. Combined with our findings that without MeCP2, regulatory elements normally low in activity in a neuronal subclass become activated and drive aberrant gene expression, these observations suggest that mCA and MeCP2 are particularly important for regulating genes that are differentially expressed between subclasses and even higher resolution neuron types.

**Fig. 4.**
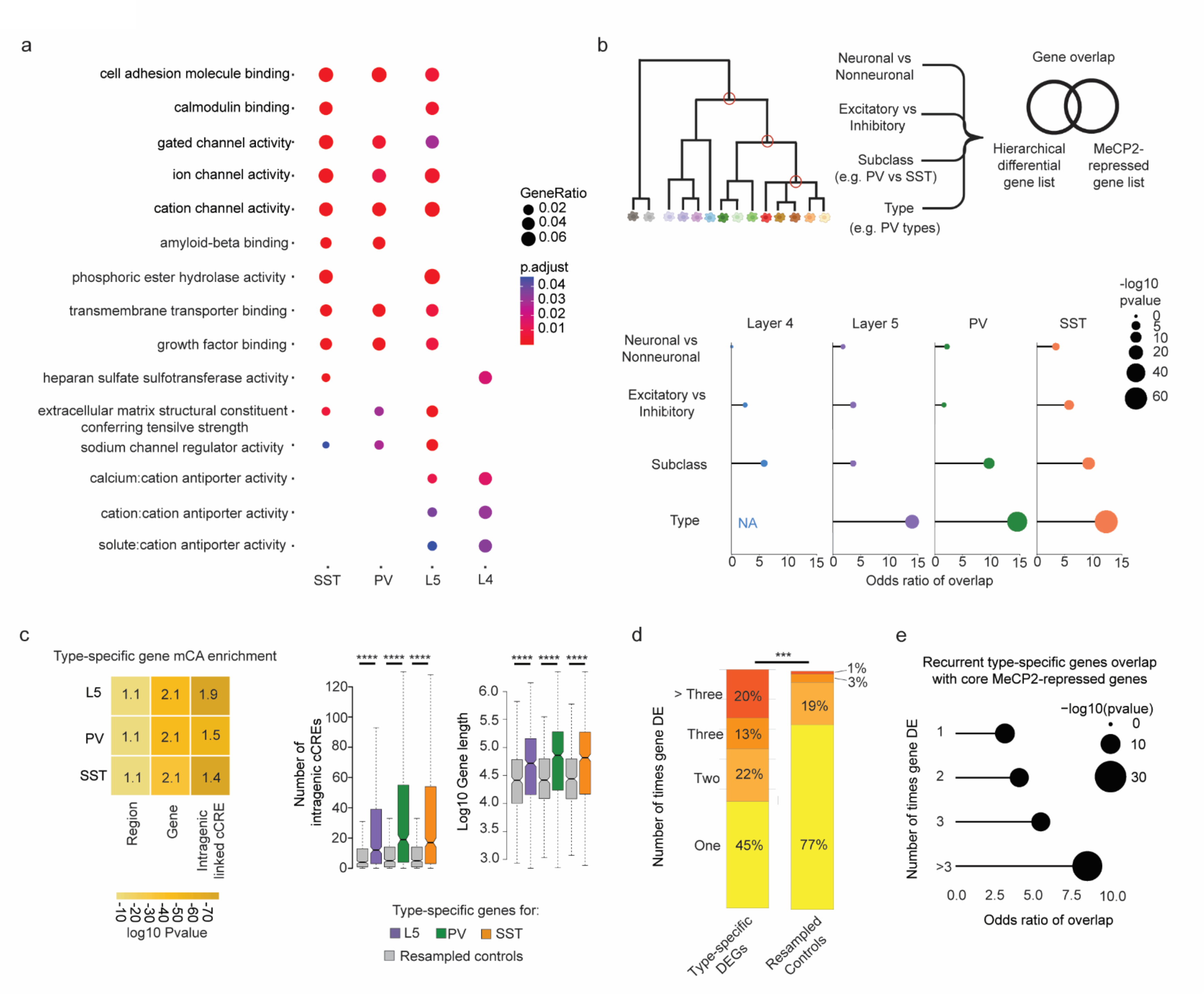
MeCP2 regulates genes that are repeatedly tuned across closely related neuronal types. **a,** Gene Ontology analysis of significantly dysregulated genes in each subclass from INTACT analysis shows enrichment for synaptic proteins, channels, and other factors important for neuronal cell type function. Top Molecular Function terms with overlap between cell types are shown. Gene ratio (percentage of total DEG in given GO term) and Benjamini-Hochberg adjusted p-value shown. **b,** Top: schematic of gene overlap analysis of MeCP2-repressed genes with gene sets distinguishing neuronal populations at different levels of hierarchical taxonomy previously generated from single cell transcriptomic data^55^. Bottom: odds ratio and significance of overlap between MeCP2-repressed genes identified in each neuron subclass and genes differentially expressed at each level of the hierarchy. P-value determined by two-sided Fisher’s exact test. NA shown for L4, because this subclass does not contain any defined types in the cellular taxonomy analyzed. **c,** Left, mCA enrichment analysis of regions, gene bodies, and intragenic-linked cCREs for genes that distinguish neuronal types within L5, PV, and SST subclasses. Enrichment relative to expression resampled control gene sets is shown numerically, significance is indicated by color. Right, quantification of number of intragenic enhancers and gene length for these neuron-type-specific genes. The center line of each boxplot is the median. Each box encloses the first and third quartile of the data. The whiskers extend to the most extreme values, excluding outliers which are outside 1.5 times the interquartile range. ****p < 0.0001 two-sided Wilcoxon rank-sum test **d,** Percentage of genes found to be differentially expressed between closely related neuron types in one or more pairwise comparisons of types within L5, PV, and SST subclasses. Values shown for true type-specific genes and for expression-matched, resampled control genes. **e,** Repeatedly tuned neuron type-specific genes significantly overlap with core MeCP2-repressed genes.

To further investigate if MeCP2 regulates genes that define neuron types, we turned to hierarchically organized single cell RNA-sequencing taxonomy from the primary visual and anterior lateral motor cortices^55^. We identified gene sets that distinguish cells at each level of the hierarchy – starting at genes that distinguish neurons from non-neuronal cells down to those that distinguish neuronal types within a subclass of neurons. We then assessed the overlap of the gene sets at each level of this hierarchy with the MeCP2-repressed genes identified in each subclass (Fig. 4b). Strikingly, we find that MeCP2-repressed genes are enriched for genes that define differences between closely related cell populations, with substantial overlap for genes that distinguish subclasses of neurons (e.g., PV vs. SST) and remarkable enrichment of genes that distinguish types of neurons at the highest resolution within subclasses (e.g., PV types) (Fig. 4b). This finding is not dependent on performing gene expression analysis with gene lists identified in specific neuronal subclasses, as we find a similar pattern of overlap with core MeCP2-repressed genes, as well as MeCP2-repressed genes identified in whole cortex analysis (Extended Data Fig. 5b). Further, enrichment of type-specific genes is also specific to MeCP2 gene regulation, as gene sets altered in other neurodevelopmental models show overlap with neuronal genes, but do not show the same preferential enrichment for genes that distinguish neuronal subclasses and types (Extended Data Fig. 5b).

We and others have shown that MeCP2 can have subthreshold effects on genes outside of those detected as significantly dysregulated^10,12,13,15^. We therefore considered the possibility that beyond overlap of gene lists, type-specific genes may be preferentially targeted by MeCP2 as a population. We assessed the characteristics of type-distinguishing genes to see if they shared features that predispose them to MeCP2 regulation. Indeed, we find that neuronal type-distinguishing genes tend to be located in regions of high mCA, are enriched for mCA in their gene bodies, and contain large numbers of intragenic cCREs that are preferential targets of MeCP2 (Fig. 4c). Thus, neuronal type-defining genes as a population have characteristics that predispose them to MeCP2 regulation, supporting the hypothesis that loss of cell-type-specific repression by MeCP2 leads to disruption of gene expression patterns that differentiate highly related, but distinct neuron types. Finely resolved types of neurons can arise within subclasses during postnatal development^2^, and have recently been shown to have distinct synaptic targets within cortical microcircuits^70^. Thus, disruption of these gene programs could contribute to circuit dysfunction in Rett syndrome and other DNA methylation pathway disorders.

While these data implicate mCA and MeCP2 in defining type-specific gene programs, they appear to contradict ample evidence of overlapping target genes for MeCP2 across cell types and brain regions. How can MeCP2 regulate highly specific gene programs but still consistently impact the same genes across brain regions and broad cell populations? One explanation for these paradoxical findings could be that the same groups of genes may be differentially expressed between closely related neuronal types found within different subclasses and brain regions (e.g., the same differentially expressed genes are tuned between two types within the PV subclass, and also between two types within the L5 subclass). Indeed, analysis of type-specific genes across the subclasses revealed that many genes are in fact differentially expressed between pairs of cell types within multiple different subclasses, and these “repeatedly tuned” genes occur at rate significantly more than would be predicted by chance (Fig. 4d). Notably, the more often a gene is found to be differentially expressed between types, the longer the gene tends to be and the higher numbers of intragenic regulatory elements the gene contains (Extended Data Fig. 6a). Furthermore, genes that are repeatedly tuned tend to be located in regions of high mCA (Extended Data Fig. 6b). Consistent with these characteristics driving regulation by MeCP2 across many cell types, repeatedly tuned genes overlap significantly with core MeCP2-repressed genes (Fig. 4e). Together, these analyses show that genes which are tuned across many closely related neuronal types have genomic characteristics (long length, many intragenic enhancers, high regional mCA) that predispose them to come under regulation by the MeCP2 pathway, and suggest that a major function of this pathway is to regulate differential gene expression between cells at the finest scale of neuronal type distinctions.

### Spatial transcriptomics reveal high-resolution gene dysregulation resulting from MeCP2 mutation

Given this new evidence implicating mCA and MeCP2 in regulating high resolution differential gene expression, we sought to extend our study to the highest level of cellular resolution. We therefore turned to Multiplexed Error Robust Fluorescence In Situ Hybridization (MERFISH)^56^, a non-amplification-based RNA quantification approach that would allow us to simultaneously define cell types, determine their location, and assess changes in gene expression for hundreds genes with single-cell spatial resolution. We designed a MERFISH gene detection panel that probed expression of 490 genes, including genes previously employed to identify cell types in single cell studies of the mouse brain^57–59^, core MeCP2-repressed genes, the MeCP2 gene itself, and additional genes detected as repeatedly tuned across neuron types. Finally, we incorporated control genes that are short and lowly methylated across cell types, and therefore likely excluded from regulation by MeCP2 (Table S2). To assess the impacts on gene expression in a highly controlled, Rett syndrome relevant paradigm, we performed our MERFISH analysis on 8-week-old *Mecp2* heterozygous null (MeCP2 KO/+) female mice that model women with Rett syndrome. Because *Mecp2* is an X-linked gene, MeCP2 KO/+ mice (and Rett syndrome patients) contain a population of cells expressing only the wild-type *Mecp2* allele and a population expressing only the deleted gene. This provides a powerful system to directly compare wild-type (WT) and knockout (KO) cells within one experiment^60^ and allowed us to quantify cell-autonomous effects of MeCP2 loss while controlling for the efficacy of *in situ* hybridization within the same analyzed brain section.

Four coronal sections containing cortex and hippocampus were isolated at ~bregma −3 mm from three independent MeCP2 KO/+ brains, and imaged on the MERSCOPE platform (Fig. 5a, Extended Data Fig. 7). After quality filtering and total count per cell normalization, the Seurat label transfer pipeline^61^ was used to map each cell to types within a compendium of cell types defined by single-cell RNA-seq of the cortex and hippocampal formation^1^ (see methods). This yielded 116,387 high-quality, pass-filter cells corresponding to 273 different excitatory, inhibitory, and non-neuronal cell types (Fig. 5b,c). Independent experiments showed robust reproducibility in transcript detection, relative gene expression, and co-clustering of cells across replicates (Extended Data Fig. 7a,b). Locations of cell types within the cortex and hippocampus showed good correspondence with previously observed distributions^1,58^ including cortical layer- and hippocampal subregion-specific localization of excitatory neurons, and cortical layer-specific distributions of inhibitory neuron types (Fig. 5b). Comparison of our bulk INTACT-RNA-seq data to expression measured by MERFISH revealed strong correlations in gene expression levels across these analysis paradigms (Extended Data Fig. 7c). Relative numbers of distinct cell types and their expression of key marker genes were also in agreement with published studies^1^ (Extended Data Fig. 7d,e).

**Fig. 5.**
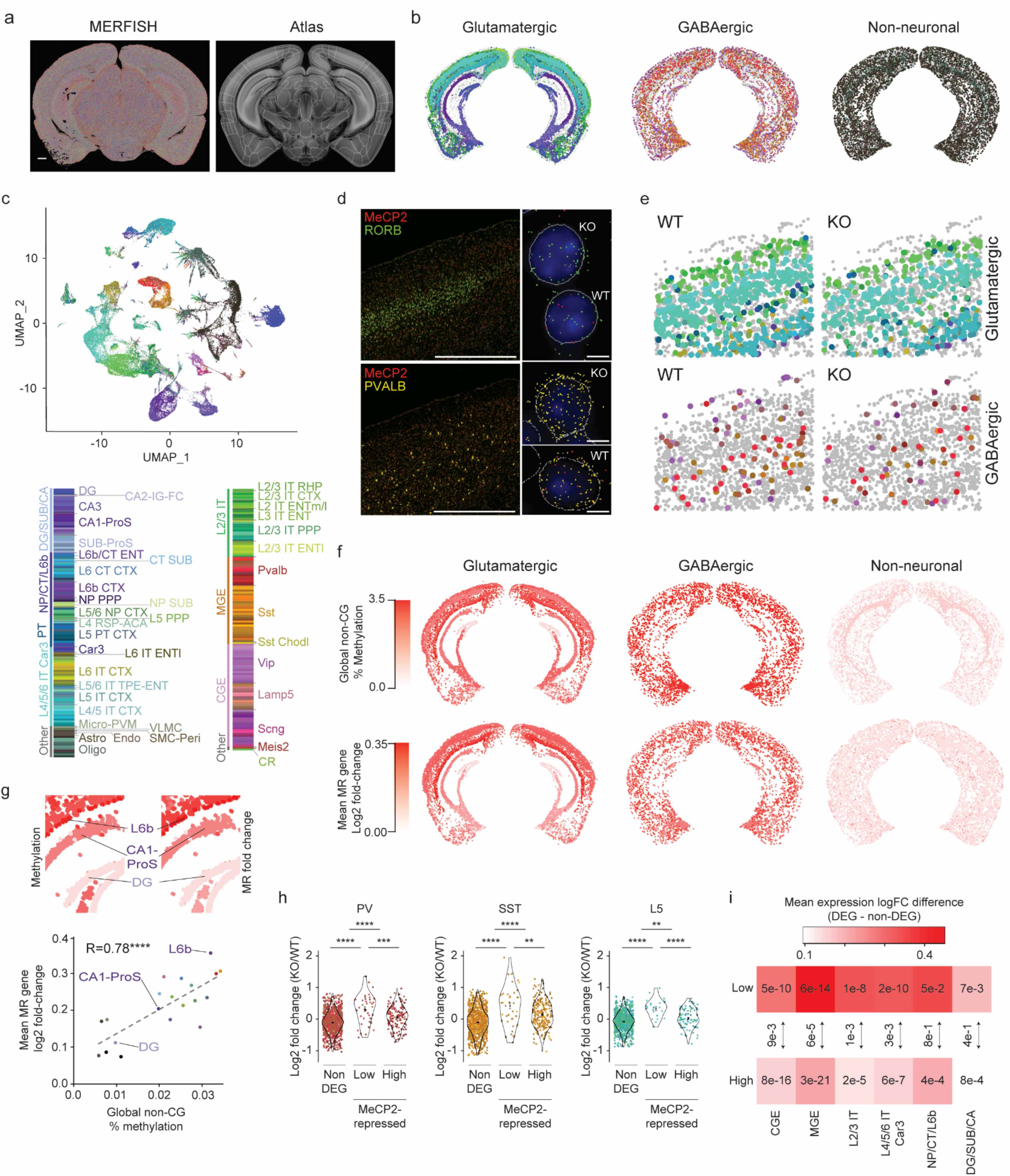
Spatial transcriptomic analysis of the Rett syndrome mouse model reveals region- and neuron type-specific gene dysregulation. **a,** Left: transcripts detected (colored dots) by MERFISH for an example MeCP2 KO/+ coronal section. Right: brain regions in the coronal plane, as defined in the Allen Mouse Brain Atlas, were identified using SHARP-Track^87,88^. Scale bar = 1 mm. **b,** Spatial map of glutamatergic, GABAergic, and non-neuronal cells in a representative MeCP2 KO/+ MERFISH experiment. Colors indicating cell types correspond to legend in **c**. **c,** Top: UMAP representation of all pass-filter cells identified in MeCP2 KO/+ MERFISH experiments (n=4 imaged sections from 3 biological replicates). Cells colored by type assigned by Seurat label transfer using the reference mouse cortical and hippocampal scRNA-seq compendium^1^. Bottom: legend showing neighborhoods, subclasses, and types of cells detected. **d,** Left: example images of cortex from MeCP2 KO/+ mice showing expression of MeCP2 (red) and L4 marker Rorb+ (top) or Pvalb (bottom) transcripts. Scale bar = 1 mm. Right: zoom views show examples of cell segmentation and transcripts detected for WT and KO transcriptotyped cells. DAPI nuclear stain is shown in blue. Scale bar = 10 µm. **e.** Spatial map of WT and KO transcriptotyped glutamatergic and GABAergic cells in MeCP2 KO/+ cortex. **f,** Top row: spatial map of glutamatergic, GABAergic, and non-neuronal subclasses, colored by global non-CG methylation level of each subclass. Bottom row: spatial map of glutamatergic, GABAergic, and non-neuronal subclasses, colored by mean log2 fold-change of core MeCP2-repressed genes in that subclass. **g,** Top: zoom-in view of glutamatergic cells shown in **f** showing global non-CG methylation levels and fold-change of core MeCP2-repressed genes in subclasses of cells. Bottom: scatter plot of mean log2 fold-change of core MeCP2-repressed genes and global non-CG methylation levels for all subclasses that can be mapped between MERFISH and single cell methylomic data^7^. **h,** Log2 fold-change of non-type-specific genes (non-DEGs), MeCP2-repressed genes that are normally “low” in each neuron type, or MeCP2-repressed genes that are normally “high” in each neuron type, for cell types found in the PV, SST, and L5 populations captured in our INTACT RNA-seq analyses. Violin plot shows fold-changes of the entire population of type-specific genes within each subclass. Points indicate fold-change of individual genes in each neuron type. Colors of points corresponds to the neuron type where the gene is normally lowly expressed; see color legend in panel **c**. **p < 0.01, ***p < 0.001, ****p < 0.0001 two-sided Wilcoxon test. The center black dot represents the mean and the error represents the standard error. Violin plots were made after excluding outliers which are outside 1.5 times the interquartile range of the data. **i,** Heatmap of mean log fold-change in gene expression difference between non-DEGs and type-specific genes “low” in a neuron type, or type-specific gene that are “high” in a neuron type in MERFISH for cell types aggregated by neighborhood. Numbers represent the p-value from the two-sided Wilcoxon rank-sum test compared to non-type-specific genes in each analysis. Arrows between boxes represent the p-value from the two-sided Wilcoxon rank-sum test comparing gene expression changes between low and high type-specific genes in each neighborhood analysis.

To interrogate the effects of loss of MeCP2 by MERFISH, cells were “transcriptotyped” by counting *Mecp2* transcripts detected by probes targeting the sequence of the mRNA deleted in the knockout allele (Fig. 5d, Extended Data Fig. 7f). This yielded a clear bimodal distribution of MeCP2 KO (0 counts) and WT (> 1 count) cells that was supported by comparison of MeCP2 count distributions to negative control probes within each MeCP2 KO/+ experiment, as well as counts of MeCP2 quantified within an entirely wild-type brain (Extended Data Fig. 7f,g). Pseudobulk Differential Gene Expression (pseudoBulkDGE)^62^ analysis of KO and WT cells corresponding to the populations captured in our INTACT RNA-seq showed robust upregulation of MeCP2-repressed genes we had identified, thus validating that our MERFISH approach could identify KO and WT cells within the same brain and detect alterations in gene expression associated with the loss of MeCP2 (Extended Data Fig. 7h). Together these findings indicate that our MERFISH approach accurately identifies cell types and allows the quantitative assessment of gene expression in these cells.

We first assessed spatial distributions of KO and WT cells in the brain to determine if loss of MeCP2 impacts the overall cellular anatomy of the brain. This analysis revealed broadly similar locations and patterns of cells between transcriptotypes (Fig. 5e and Extended Data Fig. 7i), including normal distributions of excitatory and inhibitory neuron subclasses as well as of non-neuronal cells across cortical layers. These findings are supported by previous histological analyses^63,64^ and suggest that loss of MeCP2 has limited impact on the generation or survival of cell types in the brain during early development, but instead primarily affects gene expression patterns within postmitotic cell types. We therefore focused additional analysis on changes in gene expression within the individual cell subclasses and types in the brain, assessing the overall magnitude of impact of loss of MeCP2 and the effect on type-specific gene expression.

To further explore the hypothesis that the degree of gene dysregulation in MeCP2 knockout cells is correlated with the global levels of non-CG methylation, we calculated the average changes in gene expression for core MeCP2 repressed genes within subclasses of cells and compared these effects to the global methylation levels previously detected in corresponding subclasses in single-cell methylomic studies of the cortex and hippocampus (Fig. 5f,g)^7^. This analysis revealed a clear correlation between the levels of non-CG methylation within neural cells and the upregulation of MeCP2-repressed genes (Fig. 5g). For example, limited effects were detected in non-neuronal cell populations and dentate gyrus granule neurons that contain very low levels of non-CG methylation, while there was robust dysregulation in inhibitory and excitatory populations that contain high non-CG methylation (Fig. 5f,g). Notably, these effects lead to differential impacts between hippocampal subregions (e.g., dentate gyrus vs. CA1) and cortical layers (e.g., L2/3 vs. L6b). These results extend our findings from INTACT analysis (Fig. 1), indicating that when MeCP2 is mutated in disease, global differences in mCA are likely to drive differential functional disruption across fine-resolution subregions of the brain.

We next utilized our MERFISH dataset to assess dysregulation of repeatedly tuned, type-specific gene expression upon loss of MeCP2. For this analysis, we examined MeCP2-repressed genes detected in our MERFISH gene panel that had previously been shown to exhibit differential tuning across cell types within each subclass in the cortex and hippocampus^1^. Using WT cells, we classified these genes as “low” or “high” in each type based on relative expression when compared to types within the same subclass. We then evaluated the degree of dysregulation detected for these genes in MeCP2 KO cells compared to WT in pseudoBulkDGE analysis of each type within the SST, PV, and L5 populations previously assessed by INTACT (L4 cells lacked sufficient diversity of types to be assessed). This analysis revealed a robust upregulation of “low” genes across neuron types, while “high” genes were significantly less affected (Fig. 5h). To broaden our analysis, we extended this quantification to all neuron types included within larger “neighborhoods” of subclasses defined within the cortex and hippocampus cellular taxonomy^1^. We detected similar effects across these broad families, with MeCP2-repressed genes showing upregulation in neuron types in which the gene is normally lowly expressed, but showing less upregulation in cell types in which it is normally well-expressed (Fig. 5i).

As an orthogonal approach to examine type-specific gene repression by MeCP2, we performed *in situ* hybridization by RNAScope analysis in the visual cortex of a subset of genes we identified as differentially expressed in PV neurons by INTACT-RNA-seq, but that were not well-captured by the MERFISH approach due to extremely low expression. This analysis revealed an increase in KO cells positive for joint expression of pairs of genes that are normally mutually exclusive in their expression across high-resolution PV types (Extended Data Fig. 8), further suggesting a loss of type-specific repression in MeCP2 KO cells. Together, these findings support the model in which genes that are repeatedly tuned across neuron types are often repressed by MeCP2, with this repression being strongest in neurons in which the gene is normally maintained at low levels of expression. Upon loss of MeCP2, differential expression of these genes between closely related types breaks down, likely impacting the functional specialization of types and negatively impacting circuit function.

In light of these results identifying a role for MeCP2 in type-specific gene regulation, we sought to further explore how finely-tuned gene programs may be disrupted upon loss of MeCP2. Recent high resolution transcriptomic analysis of L2/3 of the primary visual cortex (V1) has revealed distinct experience-dependent gene programs that emerge in the postnatal period between refined excitatory neuron types located in superficial, intermediate, and deep sublayers of this layer during postnatal maturation^3^. Expression of these genes is implicated in specialization of cells with distinct higher order visual area projections, and plays a role in the maturation of binocularly responsive neurons in V1. Given the fine-tuned nature of this gene regulation and the fact that it is established just before high levels of mCA and MeCP2 fully accumulate, we investigated if this spatially resolved gene program might be maintained by the MeCP2 pathway. Analysis of the spatially regulated L2/3 gene sets^3^ revealed that they are biased toward long gene length, contain many intragenic enhancers, and are enriched for mCA, suggesting that they are predisposed to regulation by MeCP2 (Extended Data Fig. 9a). Indeed, these genes significantly overlap with core MeCP2-regulated genes (Extended Data Fig. 9b).

To directly interrogate the impact of MeCP2 mutation on these genes, we quantified their expression across L2/3 of the primary visual cortex in our MERFISH data. This analysis confirmed the spatially resolved expression of these genes across wild-type excitatory neurons located in L2/3 (Fig. 6a-c,e and Extended Data Fig. 9c). Consistent with a role for MeCP2 in maintaining repression of genes that are normally lowly expressed in one cell type but more highly expressed in closely related cell types, we observed upregulation of deep sublayer genes in upper layer neurons, where these genes are normally repressed (Fig. 6b-f and Extended Data Fig. 9d). Notably, we also observed a reduction in expression of superficial sublayer-specific genes in superficially-positioned MeCP2 knockout cells. Together these effects led to an overall loss of reciprocal expression of deep- and superficial-sublayer specific gene programs in MeCP2 KO cells (Fig. 6f). Notably, superficial layer genes as a population overlap with core MeCP2-activated genes (Extended Data Fig. 9b), suggesting that in addition to loss of repression of deep layer genes, mutation of MeCP2 may consistently impact these genes through loss of cell-type-specific gene regulatory networks or other activating mechanisms.

**Fig. 6.**
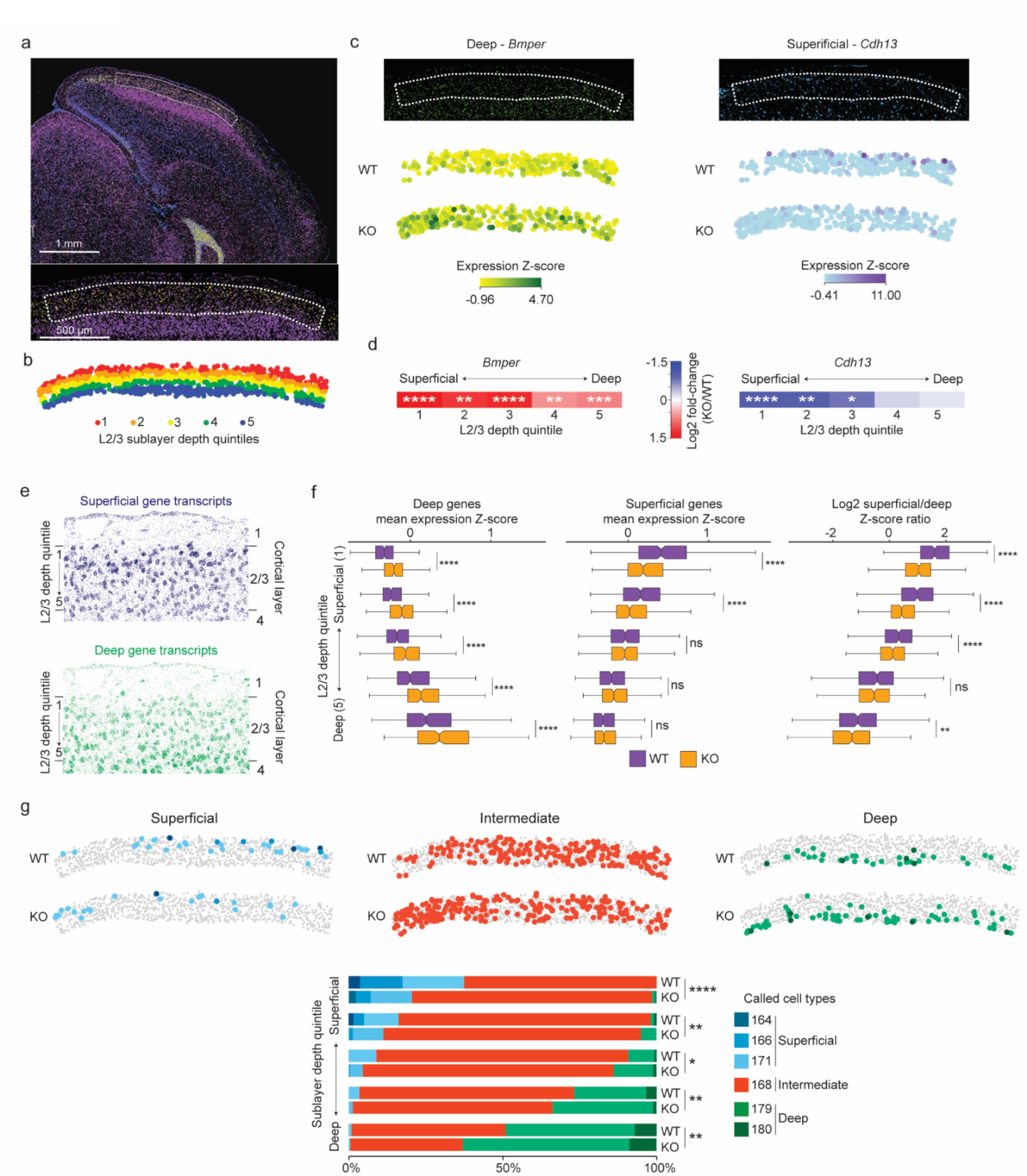
Loss of MeCP2 disrupts sublayer-resolved gene programs in layer 2/3 of the visual cortex. **a,** MERFISH image of a coronal section from an MeCP2 KO/+ brain (top) showing expression of the L2/3 marker *Ccbe1* (yellow) and the L4 marker *Rorb* (purple); blue, DAPI. Dotted line demarcates L2/3 of primary visual cortex (V1) (bottom, higher magnification) selected for further analysis. **b,** Spatial map of excitatory neurons in L2/3 for a representative MeCP2 KO/+ V1 region where each cell is colored by the quintile of sublayer depth in which it resides. **c,** Examples of MERFISH signal and transcriptotype-resolved expression in WT vs. KO excitatory neurons in L2/3 of V1 for sublayer-specific genes detected as dysregulated in the MeCP2 KO cells. Top, MERFISH signal from an example MeCP2 KO/+ brain section. Bottom, expression from cells of each transcriptotype across all excitatory neurons in L2/3 of V1. Expression is reported as the z-score of SCTransform-corrected counts of the mRNA across glutamatergic cells in L2/3 of the visual cortex. **d,** Fold-change observed in MeCP2 KO vs. WT cells across superficial to deep sublayer depth quintiles for sublayer-specific genes examined in **c**. *p. < 0.05, **p < 0.01, ***p < 0.001, ****p < 0.0001, Benjamini-Hochberg corrected p-value from pseudobulkDGE analysis. **e,** Spatial map of RNA transcripts detected in L2/3 of the MeCP2 KO/+ V1 for gene sets previously^3^ proposed to be preferentially expressed in neurons located in the superficial sublayer (top, dark blue) or deep sublayer (bottom, green) **f,** Left, and middle: boxplots of mean expression z-score of genes previously associated with deep sublayers (left) and superficial sublayers (middle) in WT and MeCP2 KO cells across sublayer depth quintiles. Right: boxplots of ratio of mean expression z-score of superficial-sublayer-associated genes to mean expression z-score of deep-sublayer-associated genes, plotted by sublayer depth quintile. The center line of each boxplot is the median. Each box encloses the first and third quartile of the data. The whiskers extend to the most extreme values, excluding outliers which are outside 1.5 times the interquartile range. **g,** Top: spatial map of WT and KO excitatory neuron IT types detected in L2/3 of the MeCP2 KO/+ V1 separated by cell types associated with superficial, intermediate, or deep gene programs^3^ (see Extended data figure 9). Bottom: stacked barplots showing proportions of neuron types in sublayer depth quintiles of L2/3 of V1, separated by transcriptotype. n=3 biological replicates for MeCP2 KO/+ MERFISH across 4 separate imaged brain sections.

Given the impact of loss of MeCP2 on spatial gene expression programs, we investigated whether these effects sufficiently impact the transcriptomes of cells to alter classification of neuron types associated with different sublayers in L2/3^3^. Analysis of excitatory neuron types assigned to L2/3 cells detected types with distinct distributions in superficial, intermediate, and deep sublayers that are similar to spatially resolved types previously described in these locations^3^ (Fig. 6g and Extended Data Fig. 9c). However, consistent with the disruption of superficial gene programs we observe, the number of neurons identified as superficial neuron types was reduced amongst KO cells, while a concomitant increase in types normally associated with deeper sublayers was detected for KO cells higher up in L2/3. Thus, loss of MeCP2 sufficiently alters spatially resolved gene expression in L2/3 to drive switching of cellular classifications in this region. These changes are likely to disrupt functional specialization of these cells associated with their projection targeting of higher-order visual areas or binocular responsivity. Together, these results indicate that loss of MeCP2 leads to disruption of sublayer-specific gene programs in L2/3 of the visual cortex, further supporting a role for the MeCP2 pathway in maintaining critical differential gene expression between closely related neuron types in the brain and implicating disruption of these gene programs in Rett syndrome.

## Discussion

The discovery that neurons contain varying levels of mCA and display highly cell-type-specific patterns of this methylation has opened questions regarding the mechanisms by which this methylation mediates gene expression in individual cell types and what its overarching function may be. Our epigenomic and spatial genomic findings in this study, together with previous results defining the patterning of mCA in the brain^12,27,41^, suggest a model in which the build-up of mCA and MeCP2 during the postnatal period establishes a repressive mechanism that helps to maintain differential expression of genes between closely related subclasses and types of neurons. In this model, genes that are repeatedly tuned between closely related types tend to be long, contain many enhancers, and are embedded in genomic regions that accumulate high levels of mCA in all neurons. These characteristics predispose these genes to repression by MeCP2 such that for neuron types in which a gene is not highly expressed in the postnatal period, high mCA levels are read out by MeCP2 to maintain low expression of the gene in that neuron type in the adult brain. In contrast, high expression of the gene within a neuron type during postnatal maturation will block accumulation of mCA across the gene and its intragenic enhancers, allowing the gene to escape repression by MeCP2 and maintain high expression in that type through the lifetime of the organism. In this way, the MeCP2 pathway facilitates the maintenance of neuron-type-specific gene expression patterns in the brain.

Identifying if there are particular neuronal populations that could be major drivers of nervous system dysfunction upon mutation of MeCP2 is essential to understanding the nature of pathology of Rett syndrome and MeCP2 duplication syndrome. Our finding that the amount of mCA within cell populations and brain substructures dictates the magnitude of gene dysregulation when MeCP2 is lost, suggests that global mCA levels are an important determinant of the impact of MeCP2 disruption across the brain. Corroborating this idea, functional experiments assessing PV and SST neurons (two high-mCA cell types) in MeCP2 mutants have shown these two populations to be significant contributors to pathologic phenotypes in Rett syndrome mouse models^35,65^. Our spatial analyses also underscore the degree to which individual subregions in the brain (e.g. dentate gyrus versus CA1 in the hippocampus) may be differentially impacted by MeCP2 loss due to differences in global mCA levels in the cell types that constitute those regions. Notably, disruption of mCA is implicated in additional disorders including Tatton-Brown Rahman syndrome and Sotos syndrome, and it is therefore likely that high-mCA cell types and brain structures will be differentially impacted in these disorders as well. As tools for cell-type-specific targeting and interventions grow^66–68^, these analyses can help direct therapeutic approaches for mCA and MeCP2 pathway disorders aimed at particularly susceptible neuronal classes and brain structures.

An open question regarding mCA is why some neuronal types have higher mCA levels than others. Our finding that mCA is read out by MeCP2 to stabilize neuronal type-specific gene expression may provide a clue as to the significance of these differences. Notably, multiple single-cell transcriptomic studies have shown that the high mCA PV and SST interneurons are comprised of numerous distinct types, while glia and excitatory neurons with very low levels of mCA (e.g., astrocytes, L4 neurons, dentate gyrus granule neurons) have few to no identifiable types^1,55,69^. It is intriguing to consider that the presence of high levels of mCA within a subclass of cells may facilitate the diversification and stabilization of high-resolution types, and that these neurons in fact can maintain their diverse transcriptomic states in part because of the robust effects of MeCP2 pathway in these cells.

Apparently conflicting findings in the field have raised the question of whether MeCP2 regulates different targets in different cell types^22–24^ or if there is shared regulation between brain regions and cell classes^13,38^. Our finding that mCA and MeCP2 facilitate repeated tuning of genes within neuron types provides an explanation for these findings: in any given pair of tissues or cell classes, a portion of the genes will have distinct methylation patterns that lead to differential regulation by MeCP2. However, MeCP2 represses a large number of repeatedly tuned genes in any subset of high-resolution neuron types found within each sample analyzed. Upregulation of these genes within these different types upon loss of MeCP2 can manifest as an overall subtle upregulation of the entire population of repeatedly tuned genes across samples. Thus, mCA and MeCP2 do impact a core set of genes across different populations of neurons, but in fact these core genes contribute to fine-scale tuning in distinct neuron types found within these populations.

Our epigenomic profiling in PV neurons indicates that MeCP2 controls enhancers in individual subclasses to regulate gene expression. This extends recent findings in whole tissue that MeCP2 blocks transcriptional initiation by inhibiting enhancer activity^10,12^. Interestingly, only half of PV MeCP2-repressed cCREs are robustly active in PV neurons under normal conditions, while the remainder show low activity in PV cells with higher activity in other cell types. Neuronal genes contain a multitude of enhancers that provide different transcriptional instructions in different types^42,71–73^. Without MeCP2 repression, the over-activation of cCREs in inappropriate neuron types appears to drive expression of non-cognate genes and have deleterious effects on that cell’s physiology.

Our MERFISH analysis of gene dysregulation in the MeCP2^KO/+^ brain demonstrates that the mCA and MeCP2 help to maintain differential gene expression between closely related neuron types, including spatially segregated gene programs in sublayers of L2/3 in the visual cortex that emerge during postnatal development. A growing body of literature indicates that neurons undergo significant transcriptional maturation during the postnatal period, with the full collection of neuronal types not developing until neurons settle into their mature positions in the cortex and refine their synaptic inputs^2,3,74–76^. Developing neurons modulate their characteristics based on local cues in the early postnatal brain to configure themselves to function in their resident circuit^74,77^, including in the visual cortex where experience is required for the emergence of distinct neuronal gene programs^3^. After this period of plasticity, neurons must stabilize their properties in order to maintain functional circuits into adulthood. This process requires fine-tuning transcriptional programs to control proper expression of genes that encode synaptic proteins critical to specialized functions. Previous studies have noted that experience can shape neuronal methylomes across subclasses^41^. Based on these previous studies and our findings here, we propose that mCA and MeCP2 accumulate in neurons during the postnatal period to stabilize high-resolution, type-specific differential gene expression after the period of plasticity and environmental interaction required to define their identity. This mechanism can thereby provide epigenetic robustness to the terminal differentiation process while still allowing a period for functional diversification driven by extrinsic input. When MeCP2 is lost in Rett syndrome neurons, fluctuations between transcriptomic states in closely related neurons may ultimately lead to circuit dysfunction. In this way, the MeCP2 pathway may have evolved, at least in part, as a mechanism to stabilize functional specialization of extraordinarily diverse neuronal types and facilitate the complexity of the mammalian brain.

## Methods

### Mice

All animal protocols were approved by the Institutional Animal Care and Use Committee and the Animal Studies Committee of Washington University in St. Louis, and in accordance with guidelines from the National Institutes of Health (NIH). Pvalb-Cre mice (B6.129P2-Pvalb^tm^^1(cre)Arbr^/J) and Sst-IRES-Cre mice (Sst^tm2.1(cre)Zjh^/J) were obtained from The Jackson Laboratory. Nr5a1-Cre mice (FVB-Tg(Nr5a1-cre)2Lowl/J) were generously shared by the Allen Brain Institute, and Rbp4-Cre mice (Tg(Rbp4-cre)KL100Gsat) were generously provided by Bernardo Sabatini (Harvard University) and Yevgenia Kozorovitskiy (Northwestern University). Each of these Cre lines were crossed with Sun1:GFP mice (B6;129-Gt(ROSA)26Sor^tm5(CAG-Sun1/sfGFP)Nat^/J) obtained from The Jackson Laboratory. MeCP2 knockout mice (B6.129P2(C)-MeCP2^tm1.Bird^/J) were obtained from The Jackson Laboratory. For PV-Cre, SST-Cre, and Nr5a1-Cre, female heterozygous MeCP2 knockout mice were crossed to Cre:Sun1:GFP mice to generate hemizygous male knockout mice and wild-type male littermates. For Rbp4-Cre, we noticed recombination in Rbp4-Cre:Sun1:GFP mice so we crossed Rbp4-Cre to MeCP2:Sun1:GFP heterozygous females. *MeCP2* overexpression mice (FVB-Tg(MECP2)3Hzo/J) were cryo-recovered from The Jackson Laboratory. Female heterozygous mice (*MeCP2*^Tg3/+^) were crossed to Pvalb-Cre:Sun1:GFP mice to generate hemizygous male transgenic mice (*MeCP2*^Tg3/y^) and wild-type male litter mates (*MeCP2*^+/y^).

### INTACT

The mouse cortex was quickly dissected in ice-cold homogenization buffer (0.25 M sucrose, 25 mM KCl, 5 mM MgCl2, 20 mM Tricine-KOH) and flash frozen in liquid nitrogen and stored at −80°C. Tissue was thawed on ice in homogenization buffer containing 1 mM DTT, 0.15 spermine, 0.5 spermidine, EDTA-free protease inhibitor, and RNasin Plus RNase Inhibitor (Promega N2611) at 60 U/mL for RNA experiments. Tissue was minced using razor blades then dounce homogenized in homogenization buffer using 5 strokes with the loose pestle and tight pestle. A 5% IGELPAL-630 solution was added and the homogenate further dounced 10 times with the tight pestle. The homogenized sample was filtered through a 40 μm strainer and underlaid with a density gradient. The sample was then slowly spun at 8,000 g on a swinging bucket rotor and the nuclei collected from the density interface. Nuclei were then isolated using GFP antibody (Fisher G13062) and Protein G Dynabeads (Invitrogen 10003D) with all immunoprecipation steps being performed in a 4°C cold room.

### Nuclear RNA-seq

RNA from SUN1-purified nuclei was extracted using RNeasy Micro Kit (Qiagen) following the manufacturer’s instructions and sequencing libraries prepared using the Nugen/Tecan Ovation SoLo RNA-Seq Library Preparation Kit. Libraries for PV samples were sequenced using Illumina HiSeq 3000 (GTAC). All other experimental libraries were NextSeq 500 (Center for Genome Sciences at Washington University).

### Chromatin immunoprecipitation protocol

INTACT isolated nuclei were input into a chromatin immunoprecipitation (ChIP) experiment following a previously described protocol^12,78^. ChIP was performed for H3K27ac (0.03 μg; Abcam ab4729). ChIP libraries were generated using Ovation Ultralow Library System V2 (NuGEN). Libraries were pooled to a final concentration of 8-10 nM and sequenced using Illumina HiSeq 3000 with GTAC to acquire 15-30 million single-end reads per sample.

### Chromatin immunoprecipitation analysis

Sequenced reads were mapped to the mm9 genome using bowtie2 alignment, and reads were extended based on library sizes and deduplicated to consolidate PCR duplicate reads. Deduplicated reads were used to quantify read density normalized by the number of reads per sample and by read length in basepairs. Bedtools coverage -counts was used to quantify ChIP-seq signal at non-promoter cCREs ^42^, defined as cCREs greater than 500 bp away from a gene’s transcription start site (TSS). EdgeR was then used to determine differential ChIP-signal across genotypes. The nominal p-values from edgeR were then combined using the Fisher method (log-sum) and were Benjamini-Hochberg corrected. Acetyl peaks with a combined q-value < 0.1, and a log2 fold-change > 0 in the KO and a log2 fold-change < 0 in the OE were called MeCP2-repressed peaks, while peaks with a combined q-value < 0.1, and a log2 fold-change < 0 in the KO and a log2 fold-change > 0 in the OE were called MeCP2-activated peaks. For plots of the log2 H3K27ac fold-change of cCREs, H3K27ac ChIP signal was calculated for 1500 bp windows centered at the middle of each cCRE before calculating the log2 H3K27ac fold-change through edgeR.

### RNA sequencing analysis

RNA sequencing analysis was performed as previously described^12^. Briefly, raw FASTQ files were trimmed with Trim Galore and rRNA sequences were filtered out with Bowtie. Remaining reads were aligned to mm9 using STAR^79^ with the default parameters. Reads mapping to multiple regions in the genome were then filtered out, and uniquely mapping reads were converted to BED files and separated into intronic and exonic reads. Finally, reads were assigned to genes using bedtools coverage -counts^80^.

For gene annotation we defined a “flattened” list of longest transcript forms for each gene, generated on Ensgene annotations and obtained from the UCSC table browser. For each gene, Ensembl IDs were matched up to MGI gene names. Then, for each unique MGI gene name, the most upstream Ensgene TSS and the most downstream TES were taken as that gene’s start and stop. Based on these Ensembl gene models, we defined TSS regions and gene bodies. Exonic reads were filtered for non- and lowly-expressed coding genes (minimum of 5 counts across samples) and then DESeq2^81^ performed using adaptive shrinkage. To enable comparisons across cell types we used RUVg to normalize data from each cell type on “in silico” defined negative control genes. These were determined using RUVg recommendations as unaffected genes (bottom 5% in significant change) in KO to WT comparisons shared across all cell types. Significantly MeCP2-repressed genes were those that had a DESeq adjusted p-value < 0.1 and log2 fold-change > 0, while MeCP2-activated genes had DESeq adjusted p-value < 0.1 and log2 fold-change < 0.

### Methylation analysis

Pseudo-bulk methylomes for L4, L5, PV, and SST cells were obtained by pooling single-cell methylation data from publicly available data^8^. The pseudo-bulk methylomes were then lifted over from mm10 to mm9. The methylation level for an element was assessed by dividing the total number of reads mapping to Cs that supported mC by the total coverage in that region, using bedtools map -o sum. Gene body methylation was calculated using the region 3 kb downstream of the TSS to the TES. Regional methylation for a gene was calculated using the region 10 kb to 210 kb upstream of the TSS and the region 200 kb downstream of its TES, removing the signal from genes overlapping these regions. In Extended Data Figure 3E, methylation levels were calculated for topologically associating domains (TADs). The Arrowhead algorithm was used to identify these TADs from Knight-Ruiz normalized contact matrices as previously described^82^ from mouse cortical neurons^83^. TAD/cCRE methylation correlations were calculated by first intersecting TADs with cCREs. For a given TAD-cCRE pair, the TAD methylation signal was calculated by subtracting the methylation signal of the cCRE from the methylation across the entire TAD region^12,27^.

### Candidate cis-regulatory elements (cCREs)

Candidate cis-regulatory elements (cCREs) and their linkages to genes were identified through chromatin co-accessibility analysis and RNA expression correlation analysis by the BRAIN Initiative Cell Census Network (BICCN)^42^. The coordinates of these cCREs were lifted over from mm10 to mm9 using the UCSC LiftOver tool.

cCREs most robustly regulated by MeCP2 were identified by combining analysis of H3K27ac ChIP-seq signal in cCREs in MeCP2 KO and MeCP2 OE PV nuclei. Nominal p-values and fold-changes were calculated for the cCREs using edgeR. The p-values were combined using the Fisher method (log-sum) and were Benjamini-Hochberg corrected. cCREs with a combined adjusted p-value ≤ 0.1, log2 fold-change > 0 in the MeCP2 KO, and log2 fold-change < 0 in the MeCP2 OE were considered MeCP2-repressed cCREs. cCREs with a combined adjusted p-value ≤ 0.1, log2 fold-change < 0 in the MeCP2 KO, and log2 fold-change > 0 in the MeCP2 OE were considered MeCP2-activated cCREs.

### Motif analysis

Transcription factor motif enrichment analysis was performed using HOMER^47^ on cCREs using the following parameters: findMotifsGenome.pl input.bed mm10 output -size 200 -len 8.

### Controlled resampling

A similar resampling approach was used as previously described^12^. Briefly, for every entry in a sample set (e.g., MeCP2-repressed genes), an entry in the control set (e.g., all other genes) with a similar desired characteristic (e.g., expression) was selected, generating a control set of the same size and variable distribution as the sample set.

### Tissue preparation for RNAScope

MeCP2^KO/+^ female mice at 8 weeks of age were perfused with ice-cold saline. The brain was removed and placed in an embedding mold (Peel-A-Way, Sigma-Aldrich) filled with a cryoprotective embedding medium (O.C.T. compound, Fisher Scientific), which was flash frozen in an isopentane bath that was pre-cooled in liquid nitrogen. Tissue was stored at −80°C for up to 3 months. The night before slicing, the brain was placed in a −20°C freezer and allowed to equilibrate to −20°C. The day of slicing, it was placed in a pre-cooled cryostat. All chambers and tools were cleaned with RNase Away (Thermo Scientific) and 70% ethanol. Slices were cut coronally at 12-14 μm with the mouse visual cortex collected using Allen Brain Atlas Mouse P56 Coronal as reference. Slides were stored at −80°C in a slide box wrapped in plastic and used within two weeks.

### RNAScope *in situ* hybridization

Prior to starting the experiment, all tools and surfaces were cleaned with RNase Away. Standard protocol from ACDBio for fresh-frozen tissue was performed. Brain slices were fixed in 4% PFA, dehydrated using increasing ethanol concentrations, and then treated with Protease IV. Probes were warmed to 40°C and then mixed and added to the tissue for 2 hours at 40°C in an RNAScope hybridization oven. Amplification reagents were then applied followed by Round 1 fluorescent probes (T1-T4) and DAPI. Imaging was performed for the first round and then the slide inserted into 4X saline sodium citrate (SSC) for at least an hour until the coverslip could easily slide off the slide, with great care taken to not damage the tissue. 10% cleaving solution was then applied to cleave Round 1 fluorophores. After washing, Round 2 fluorophores (T5-T8) were added and the sample imaged. This process was repeated to image the final four probes T9-T12. Imaging was performed on Zeiss AxioImager Z2 using 40X objective. 10-12 z stacks with 1 um spacing were obtained. Images were analyzed using FIJI with cell outlined using DAPI signal. Probe images were then manually thresholded to remove background and quantification of puncta was performed. Downstream analysis was performed using custom R scripts.

### Tissue preparation for MERFISH

To obtain fresh frozen tissue, 8–10-week-old MeCP2^KO/+^ female mice were anesthetized with ketamine/xylazine and perfused with 1x phosphate buffer saline, pH 7.4 (Gibco). The brain was quickly extracted from the cranium, placed in an embedding mold (Peel-A-Way, Sigma-Aldrich) containing O.C.T., and flash-frozen by placing the mold over an isopentane bath chilled with liquid nitrogen. The frozen O.C.T. compound block containing the brain was stored in −80°C until cryosectioning.

10 μm-thick coronal tissue sections were prepared from the O.C.T. compound-embedded brain using a cryostat (Leica CM1860). Tissue sections were mounted onto specialized MERSCOPE slides (Vizgen, PN 20400001), fixed with 4% paraformaldehyde following the Vizgen protocol and maintained in 70% ethanol in a sealed petri dish until gene probe hybridization.

### MERFISH probe hybridization

Hybridization of the tissue to gene probes for MERFISH was performed according to instructions provided by Vizgen. Briefly, after aspiration of the 70% ethanol, the 10 μm-thick tissue section mounted onto the MERSCOPE slide was washed with Sample Prep Wash Buffer (Vizgen, PN 20300001) and Formamide Wash Buffer (Vizgen, PN 20300002) while placed in the petri dish, and incubated at 37°C for 30 min in a benchtop incubator (INCU-Line, VWR). The Formamide Wash Buffer was then aspirated, and 50 μL of a custom-designed MERSCOPE gene panel mix was delivered onto the tissue section. A 2 × 2 cm piece of Parafilm M (Fisher Scientific) was carefully placed onto the tissue so that the gene panel mix was evenly spread across the tissue and no bubbles were introduced between the tissue and the film. The petri dish was closed with a lid and sealed with Parafilm M, sterilized with 70% ethanol, and placed in a humidified benchtop incubator (INCU-Line, VWR) set at 37°C for 36–48 h.

### Gel embedding and tissue clearing

After gene probe hybridization, the tissue was washed twice with Formamide Wash Buffer and incubated at 47°C for 30 min after each wash, followed by an incubation in Sample Prep Wash Buffer for at least 2 min. A gel embedding solution was prepared by combining 5 mL Gel Embedding Premix (Vizgen, PN 20300004), 10% w/v ammonium persulfate solution (25 μL), and 2.5 μL N,N,N’,N’-tetramethylethylenediamine. 50 μL of the gel embedding solution was pipetted onto the tissue, and a 20-mm coverslip (Deckgläser) was slowly placed on top of the solution after cleaning the coverslip with RNAse AWAY (Molecular BioProducts) and 70% ethanol and coating the facing-down side of the coverslip with Gel Slick Solution (Lonza). The tissue was incubated at room temperature for 1.5 h to allow the gel to polymerize. After polymerization of the gel, the tissue was incubated in 5 mL of a clearing solution comprising Clearing Premix (Vizgen, PN 20300003) and 50 μL Proteinase K (New England BioLabs) at 37°C in a humidified benchtop incubator until the tissue was cleared.

### MERFISH imaging

After clearing, the tissue was photobleached for 3 h in a MERSCOPE photobleacher for background autofluorescence quenching. The gel-embedded tissue was then washed with Sample Prep Wash Buffer (Vizgen, PN 20300001) and incubated in Vizgen’s DAPI and PolyT Staining Reagent (PN 20300021) for 15 min, followed by consecutive washes with Formamide Wash Buffer and Sample Prep Wash Buffer. An imaging activation mix was prepared by adding RNAse inhibitor (100 μL, New England BioLabs) to 250 μL Imaging Buffer Activator (VIZGEN, PN 203000015). The imaging activation mix was carefully added to a thawed MERSCOPE Imaging Cartridge via the Cartridge Activation Port, and the fluidics system of the MERSCOPE instrument was primed according to the manufacturer’s instructions. The MERSCOPE slide with the gel-embedded, hybridized tissue was then attached to the MERSCOPE flow chamber following the manufacturer’s instructions. High-magnification imaging of the tissue was performed according to on-screen instructions following selection of the applicable gene panel-specific MERSCOPE codebook, acquiring a 10x mosaic DAPI image, and selection of the imaging area.

### MERFISH data pre-processing

Cell segmentation was performed through the Watershed algorithm, as previously described^84^. DAPI images were used as the seeds and PolyT signals were used to identify cell boundaries. To remove cell segmentation artifacts in our MERFISH data, we filtered out cells with volumes less than 100 μm^3^ or more than three times the median volume of cells in each experiment, in accordance with previous work^58^. We further selected for high quality cells by removing those with less than 300 total RNA counts. To focus our analyses on the cortex and hippocampus, we used MERSCOPE’s data visualization software Vizualizer to identify cells located in those regions and filter our remaining cells from our analysis.

The R package Seurat^61^ was used to further process and analyze our MERFISH data. The cell-by-gene matrix of cortical and hippocampal cells in each MERFISH experiment was converted into a Seurat object. RNA counts were normalized using the SCTransform method^85^. To facilitate joint analyses of all our MeCP2 KO/+ MERFISH experiments, we integrated the MeCP2 KO/+ Seurat objects together using Seurat’s FindIntegrationAnchors and IntegrateData functions. UMAP analysis^86^ was performed on the integrated Seurat object for visualization of cell clusters across all our MeCP2 KO/+ experiments.

### MERFISH cell type classification

Cell types in our MERFISH experiments were identified using a single-cell RNA-seq dataset of the mouse cortex and hippocampus^1^ as a reference dataset. The cell-by-gene matrix of this reference dataset was transformed into a Seurat object, after which cells with the fewest number of detected genes (<10^th^ percentile) or the greatest number of detected genes (>90^th^ percentile) were filtered out. The Seurat object was normalized by SCTransform^85^. The Seurat object was then input into Seurat’s label transfer functions as the reference dataset and our MERFISH Seurat objects were input as query datasets, allowing us to label the neuron types in our MERFISH experiments by projecting the PCA structure of the reference dataset onto the query dataset. Broader levels of classification (e.g. subclass, neighborhood, class) were determined by matching the type label to associated broader classification labels from the scRNA-seq reference dataset’s metadata. To focus on cells that Seurat labeled more confidently, we selected cells with Seurat prediction scores greater than 0.2.

### MERFISH Transcriptotyping

We transcriptotyped cells (i.e., identified cells as MeCP2 KO or MeCP2 WT) by examining the distribution of *Mecp2* SCTransform-corrected counts in each MERFISH MeCP2 KO/+ experiment, reasoning that WT and MeCP2 KO cells would show different distributions. We searched for an *Mecp2* count cutoff that would allow us identify a comparable number of MeCP2 KO and WT cells. Furthermore, to increase our confidence in assignment of WT cells, we used a *Mecp2* count cutoff that is higher than the counts of negative control blank probes in 99% of all cells, as blank probes do not target RNA. We also examined the distribution of *Mecp2* counts in an independent MERFISH experiment carried out on wild-type brain to see if we would capture most WT cells using our *Mecp2* count threshold defined in our MeCP2 KO/+ MERFISH experiments. Through this analysis, we labeled cells with less than 1 SCTransform-corrected count as MeCP2 KO and cells with greater than 1 SCTransform-corrected count as WT (See Extended Data Fig. 7f).

### MERFISH type-specific gene expression analyses

To quantify fold differences in gene expression between WT and MeCP2 KO cells, we summed raw RNA counts of each gene across all cells of each type or subclass using the pseudobulkDGE function of the R package scran^62^. Experiment ID was used as a blocking variable to account for differences in RNA counts between experiments.

To investigate cell-type-specific gene expression responses to MeCP2 disruption, we selected genes previously identified as differentially expressed between neuron types at the highest level of resolution in cortical and hippocampal cells^1^. We identified a gene as “high” in a type if it was more highly expressed in that type relative to another type within the same subclass and if it was more highly expressed in that type than the median expression of all genes in that type’s subclass. Conversely, we identified a gene as “low” in a type if it was less expressed in that type relative to another type within the same subclass and if it was less highly expressed in that type than the median expression of all genes in that type’s subclass. We plotted the gene expression log-fold-changes of these genes to determine whether there is type-specific de-repression of genes (Fig. 6h). For analysis of neighborhoods (Fig. 6i), this analysis was performed for all types within each subclass and then all type-specific genes within subclasses found in each neighborhood were analyzed in aggregate.

### MERFISH analysis of visual cortex

To analyze cells in primary visual cortex and interrogate sublayer gene expression patterns, the primary visual cortex was first identified by using the SHARP-Track image analysis package^87,88^ to register the MERFISH imaged sections to the Allen Common Coordinate Framework^89^. For further analysis of L2/3, this region was defined by using (*Ccbe1*) and L4 (*Rorb*) marker gene expression together with nuclear density as guides to dissect layers. To divide cells in L2/3 of V1 into sublayer depth quintiles, we first calculated a normalized depth value for each cell: in each experiment, we identified a reference set of cells on the upper boundary of L2/3 and query set of MeCP2 WT or KO cells in L2/3 of V1. For each query cell, we then calculated the minimal Euclidean distance between the center of the query cell and the centers of all reference cells. The normalized depth value for each query cell was then calculated by dividing the query cell’s minimal Euclidean distance by the greatest of these minimal Euclidean distances among all query cells, such that the shallowest query cell would have a normalized depth value of 0 and the deepest query cell would have a normalized depth value of 1. We then divided the cells into quintiles of sublayer depth based on their normalized depth values and carried out quantification of gene expression changes by cell type mapping and pseudobulkDGE analysis as described above.

## Data Availability

All genomic data generated in this study have been uploaded to the NCBI GEO archive GSE237089.

## Code Availability

Code will be made available upon request.

## Acknowledgements

We thank members of the Gabel lab, as well as J. Edwards for providing support and feedback on the manuscript. Next-Generation-Sequencing was carried out through the Genome Technology Access Center at the McDonnell Genome Institute and The Edison Family Center for Genome Sciences and Systems Biology at Washington University in St. Louis. We thank M. Watson and J. Snider for MERFISH imaging support. This work was supported by NIH NICHD 5F30HD102147-02 to J.R.M., NIH NICHD 1F30HD110156-01 to N.H., NINDS R01NS04102 to H.W.G., and NIMH R01MH117405 to H.W.G.

---

**Extended Data Fig. 1.**
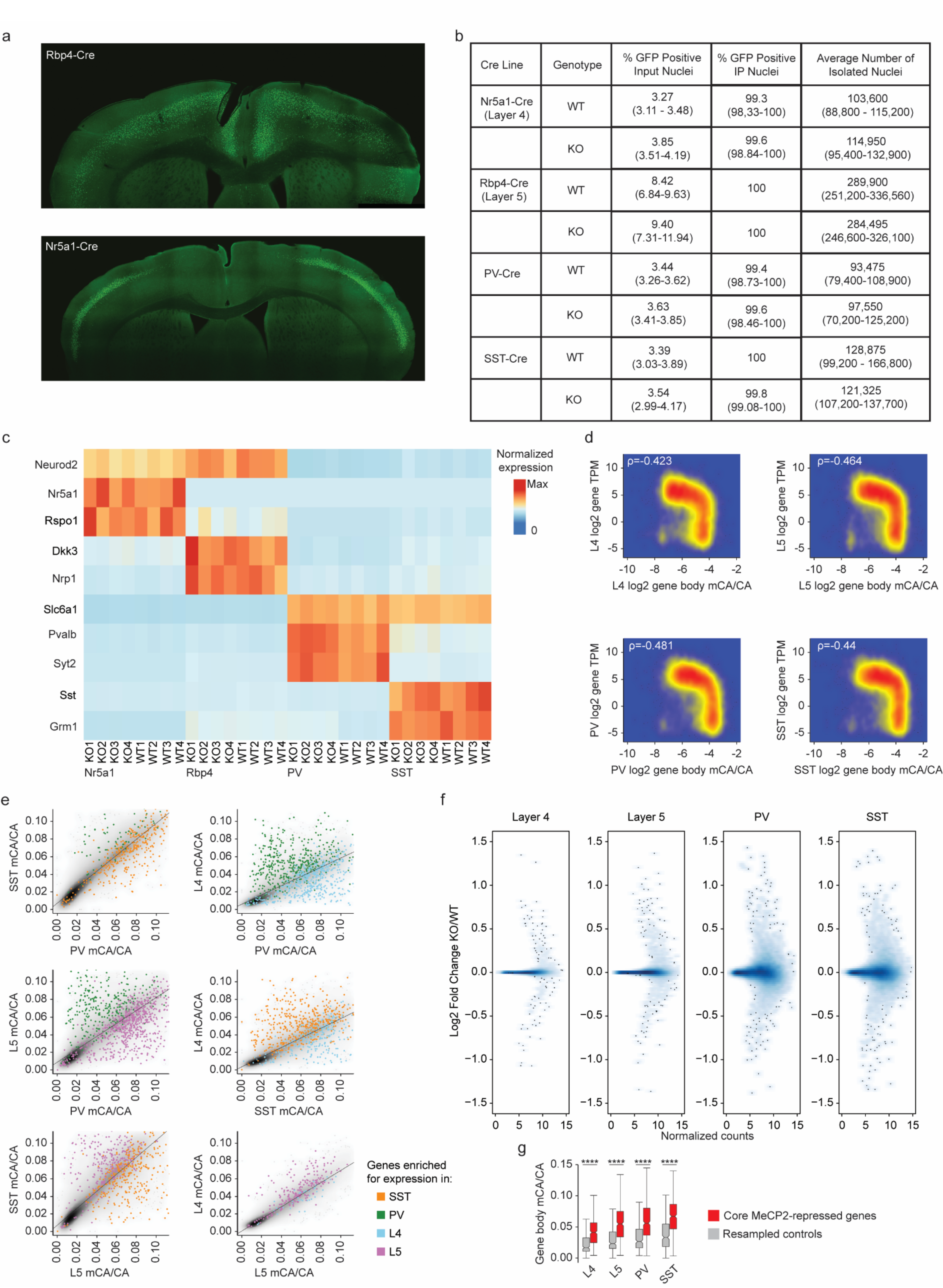
Gene expression and methylation in INTACT-isolated PV, SST, L4, and L5 neurons. **a,** Representative images of Rbp4-Cre;SUN1:GFP labeling of L5 excitatory neurons and Nr5a1-Cre;SUN1:GFP labeling of L4 excitatory neurons. **b,** Summary statistics of INTACT experiments organized by Cre-line and genotype (no significant differences found between genotypes). **c,** Heatmap of marker gene expression in RNA sequencing data from each subclass profiled. **d,** Log2 gene snmc-seq mCA/CA vs log2 gene TPMs for L4, L5, SST, and PV cells. ρ = Spearman’s correlation coefficient. **e,** Pairwise comparisons of gene body mCA/CA across cell types. Genes enriched for expression >5 fold in one cell type over another are colored according to the cell type where they are highly expressed. **f,** Log2 fold-change (MeCP2 KO/WT) of gene expression in L4, L5, SST, and PV cells. The y-axis is normalized counts of genes from DESeq2. **g,** Gene body mCA level of core MeCP2-repressed genes in each cell type compared to expression-matched, resampled control genes. ****p < 0.0001 two-sided Wilcoxon rank-sum test. The center line of each boxplot is the median. Each box encloses the first and third quartile of the data. The whiskers extend to the most extreme values, excluding outliers which are outside 1.5 times the interquartile range. n=4 biological replicates for PV, SST, L4, and L5 RNA-seq.

**Extended Data Fig. 2.**
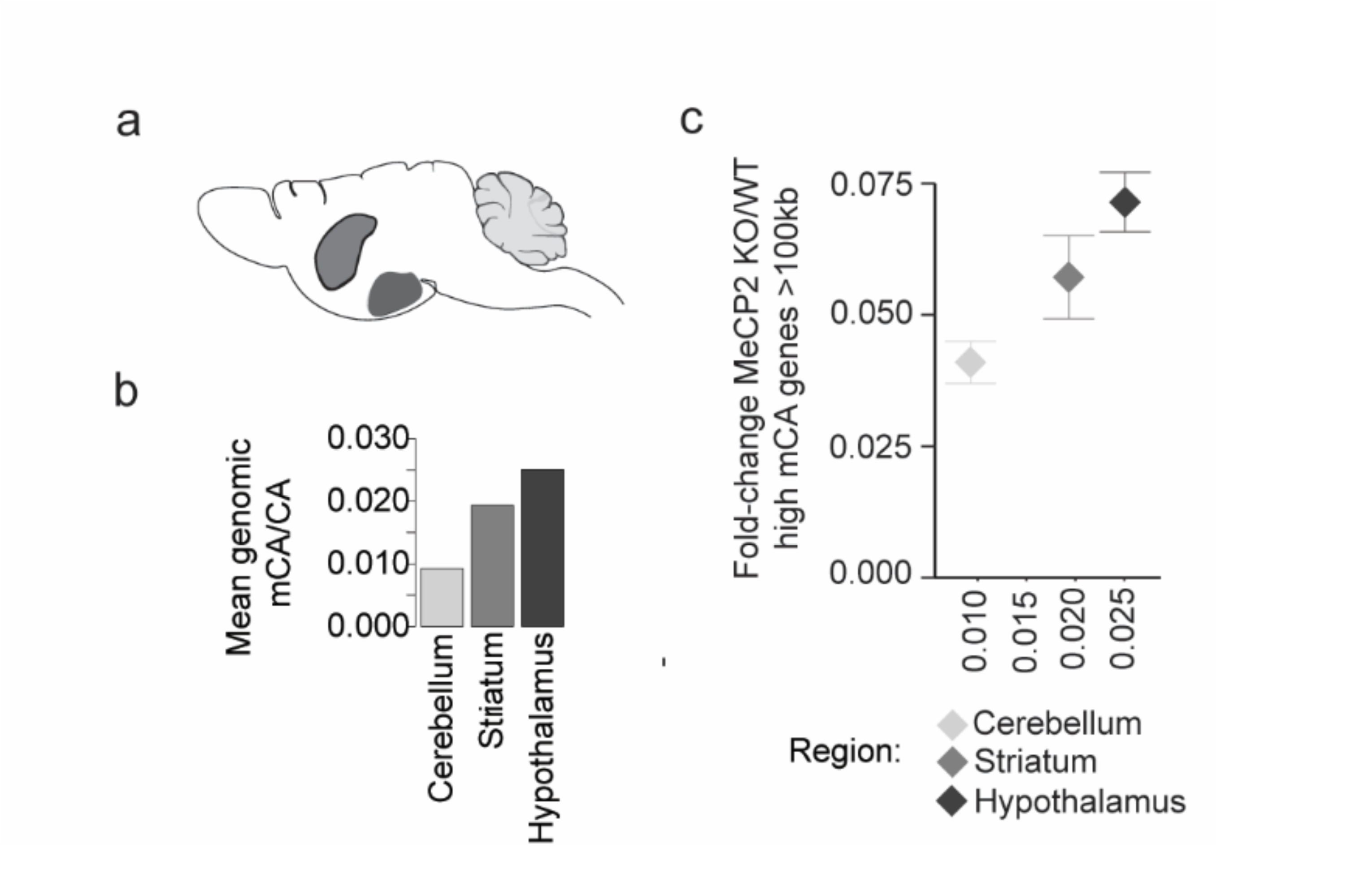
Analysis of global mCA levels and gene dysregulation in different brain regions of MeCP2 KO mice. **a,** Depiction of the three brain regions analyzed: cerebellum, striatum, and hypothalamus. **b,** Global mCA/CA levels across the three brain regions analyzed. **c,** Scatter plot of global mCA levels in each brain region and average fold-change in mRNA expression for long (greater than 100 kb), high mCA (top decile of gene body mCA) genes in the MeCP2 KO vs. WT. n=1 biological replicate for cerebellum, striatum, and hypothalamus methylation analysis.

**Extended Data Fig. 3.**
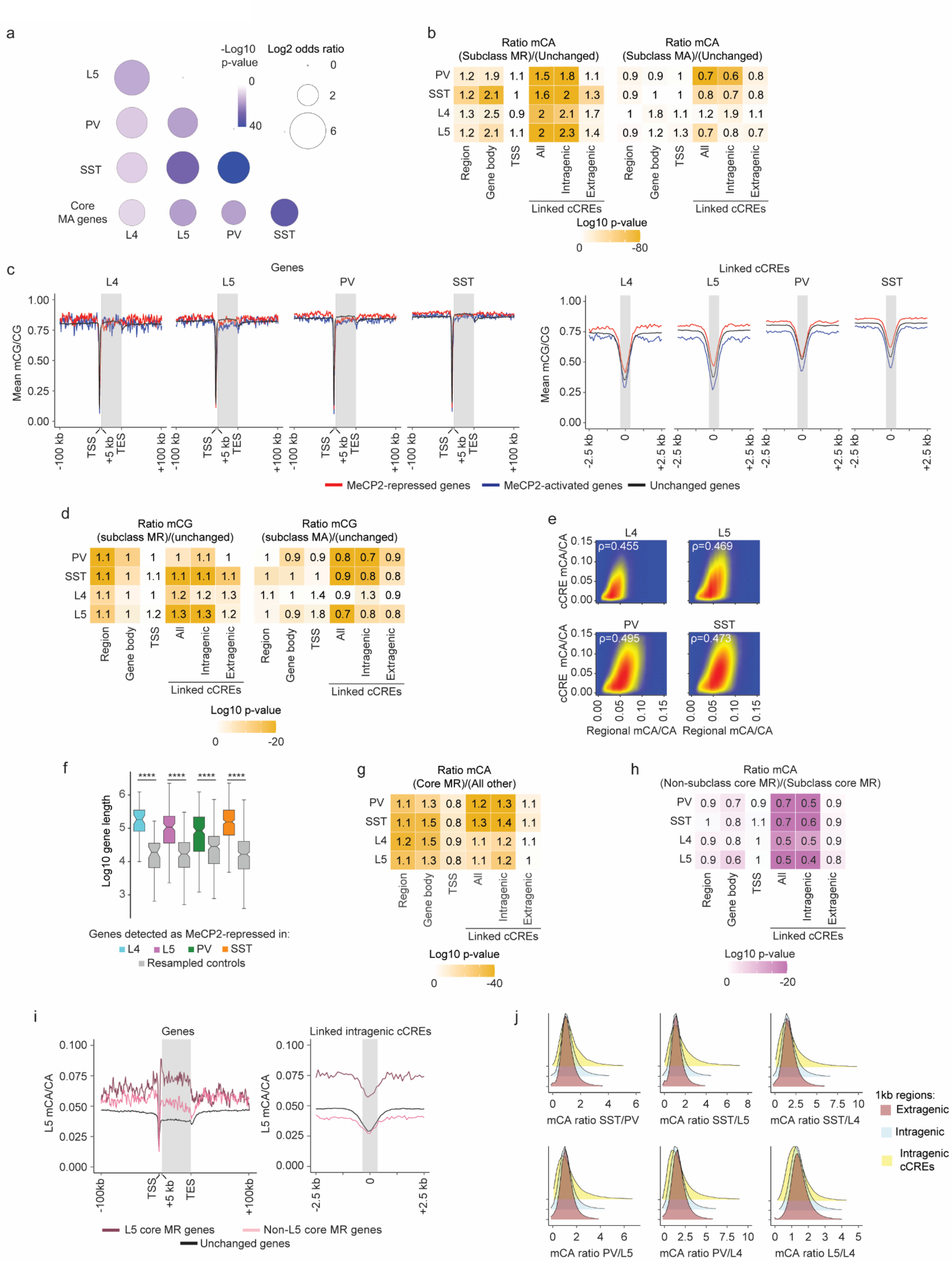
Methylation patterns at MeCP2-regulated genes in PV, SST, L4, and L5 neurons. **a,** Significance of overlap of MeCP2-activated genes from each cell type and core MeCP2-activated genes from multiple datasets. −Log10 p-value is calculated from two-sided Fisher’s exact test. **b,** Heatmap of mCA/CA enrichment in regions, gene bodies, and linked cCREs of cell-type MR genes or cell-type MA genes over those of unchanged genes, colored by the log10 two-sided Wilcoxon rank-sum p-value. Numbers in the tiles represent the ratio of median methylation level of elements associated with subclass MR or MA genes to the median methylation level of elements associated with unchanged genes. **c,** Left: Aggregate mCG/CG levels for MeCP2-regulated genes in L4, L5, SST, and PV neurons. Mean mCG/CG is reported for 1 kb bins. “Metagene’’ refers to 50 equally sized bins within gene bodies. Right: aggregate mCG levels centered at cCREs linked to MeCP2-regulated genes in L4, L5, SST, and PV neurons. Mean mCG/CG is reported for 100 bp bins. Gray rectangle = 700 bp, ~ median length of all cCREs. **d,** Heatmap of mCG/CG enrichment in regions, gene bodies, and linked cCREs of cell-type MR genes or cell-type MA genes over those of unchanged genes, colored by the log10 two-sided Wilcoxon rank-sum p-value. Numbers in the tiles represent the ratio of median methylation level of elements associated with cell-type MR or MA genes to the median methylation level of elements associated with unchanged genes. **e,** Scatter plot showing regional mCA/CA levels and cCRE mCA/CA levels for all cCREs in the genome in each cell type. This neuron subclass specific analysis shows that cCREs mCA is correlated with the mCA set-point for large-scale genomic regions, as previously shown for whole cortex^12^. Here, regions are defined as topologically associated domains (TADs) identified in the cortex^83^. ρ = Spearman’s correlation coefficient. **f,** Log 10 gene length of genes MeCP2-repressed in L4, L5, PV, and SST genes. The gray box next to each subclass represents the expression-matched genes resampled from that subclass’s list of unchanged genes. The center line of each boxplot is the median. Each box encloses the first and third quartile of the data. The whiskers extend to the most extreme values, excluding outliers which are outside 1.5 times the interquartile range. **g,** Heatmap of mCA/CA enrichment in regions, gene bodies, and linked cCREs core MR genes over those of all other genes, colored by the log10 Wilcoxon rank-sum p-value. Numbers in the tiles represent the ratio of median mCA/CA of elements associated with core MR genes to the median mCA/CA of elements associated with all other genes. **h,** Heatmap of mCA/CA enrichment in regions, gene bodies, and linked cCREs of non-subclass core MR genes over those of subclass core MR genes, colored by the log10 two-sided Wilcoxon rank-sum p-value. Numbers in the tiles represent the ratio of median mCA/CA of elements associated with subclass MR genes to the median mCA/CA of elements associated with other-subclass MR genes. **i,** Aggregate mCA/CA levels at gene bodies (left) and linked cCREs (right) of L5 core MR genes, non-L5 core MR genes, and unchanged genes. **j,** Density plots of pairwise mCA/CA ratios between cell types in 1 kb extragenic regions, intragenic regions, and regions centered at intragenic cCREs. DNA methylation data were compiled from previously published single-cell methylomic analysis^7^, *see methods*. RNA-seq data are from INTACT analysis described in **Figure 1**, n=4 biological replicates per genotype per cell type.

**Extended Data Fig. 4.**
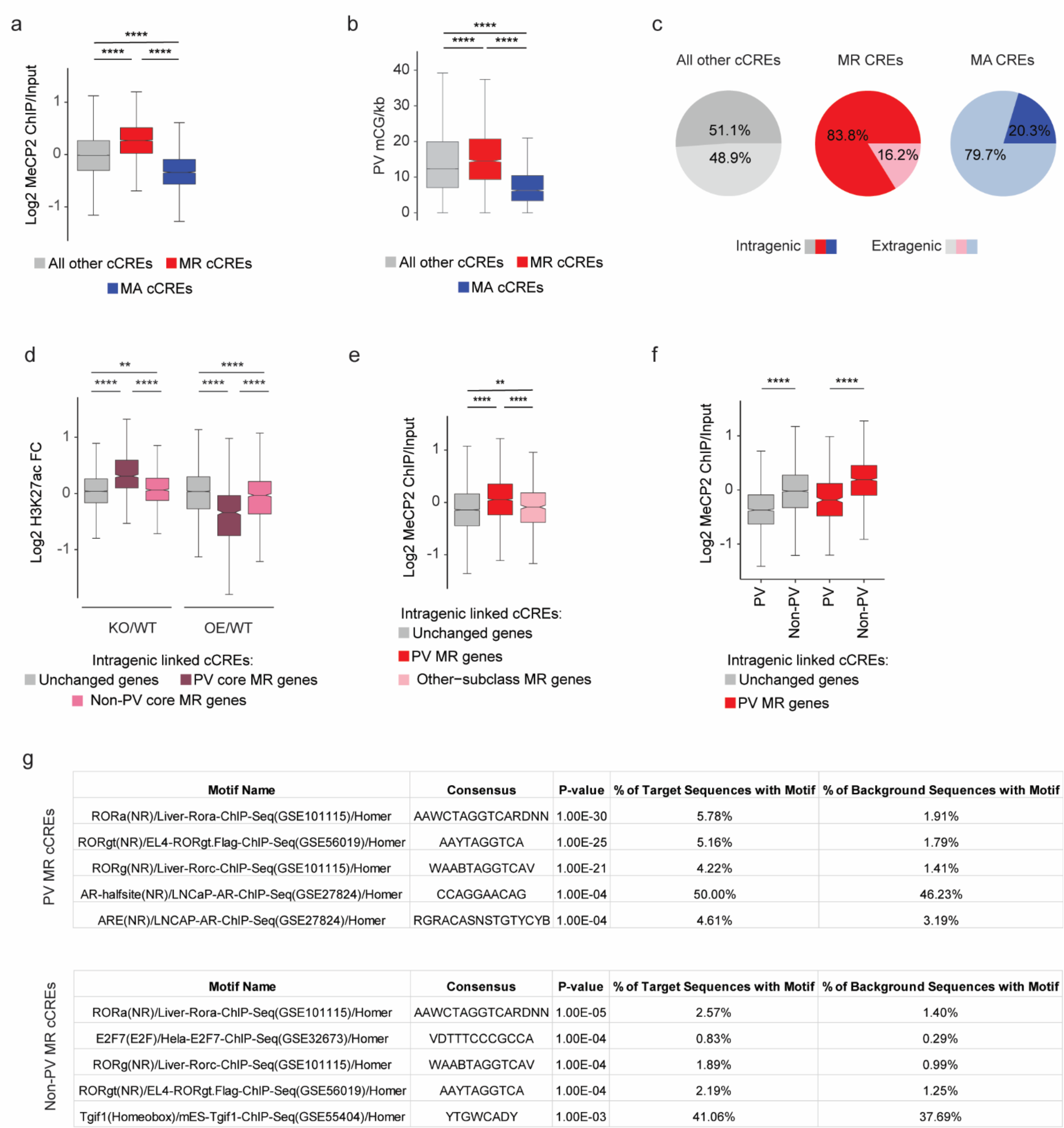
Distribution of epigenomic signals at MeCP2-regulated cCREs in PV interneurons. **a,** Log2 input-normalized MeCP2 ChIP signal at MeCP2-repressed (MR) and MeCP2-activated (MA) cCREs in PV cells. ****p < 0.0001 two-sided Wilcoxon rank-sum test. **b,** Boxplot of PV mCG/kb in MeCP2-regulated cCREs. ****p < 0.0001 two-sided Wilcoxon rank-sum test. **c,** Genic distributions of MeCP2-regulated cCREs showing enrichment of MR cCREs to be intragenic. **d,** Log2 H3K27ac ChIP fold-change (MeCP2 mutant/wild-type) in cCREs inside and linked to PV core MR, non-PV core MR, or unchanged genes. **p < 0.01, ****p < 0.0001 two-sided Wilcoxon rank-sum test. **e,** Log2 input-normalized MeCP2 ChIP-seq signal in cCREs inside and linked to PV MR genes, other-subclass MR genes, or unchanged genes. **p < 0.01, ****p < 0.0001 two-sided Wilcoxon rank-sum test. **f,** Log2 input-normalized MeCP2 ChIP-seq signal in PV cCREs and non-PV cCREs inside and linked to PV MR genes or unchanged genes. ****p < 0.0001 two-sided Wilcoxon rank-sum test. **g,** HOMER output showing ROR family motif enrichment in PV and non-PV cCREs. n=3 biological replicates for PV WT, MeCP2 KO, and MeCP2 OE H3K27ac ChIP-seq. In all boxplots, the center line of each boxplot is the median. Each box encloses the first and third quartile of the data. The whiskers extend to the most extreme values, excluding outliers which are outside 1.5 times the interquartile range.

**Extended Data Fig. 5.**
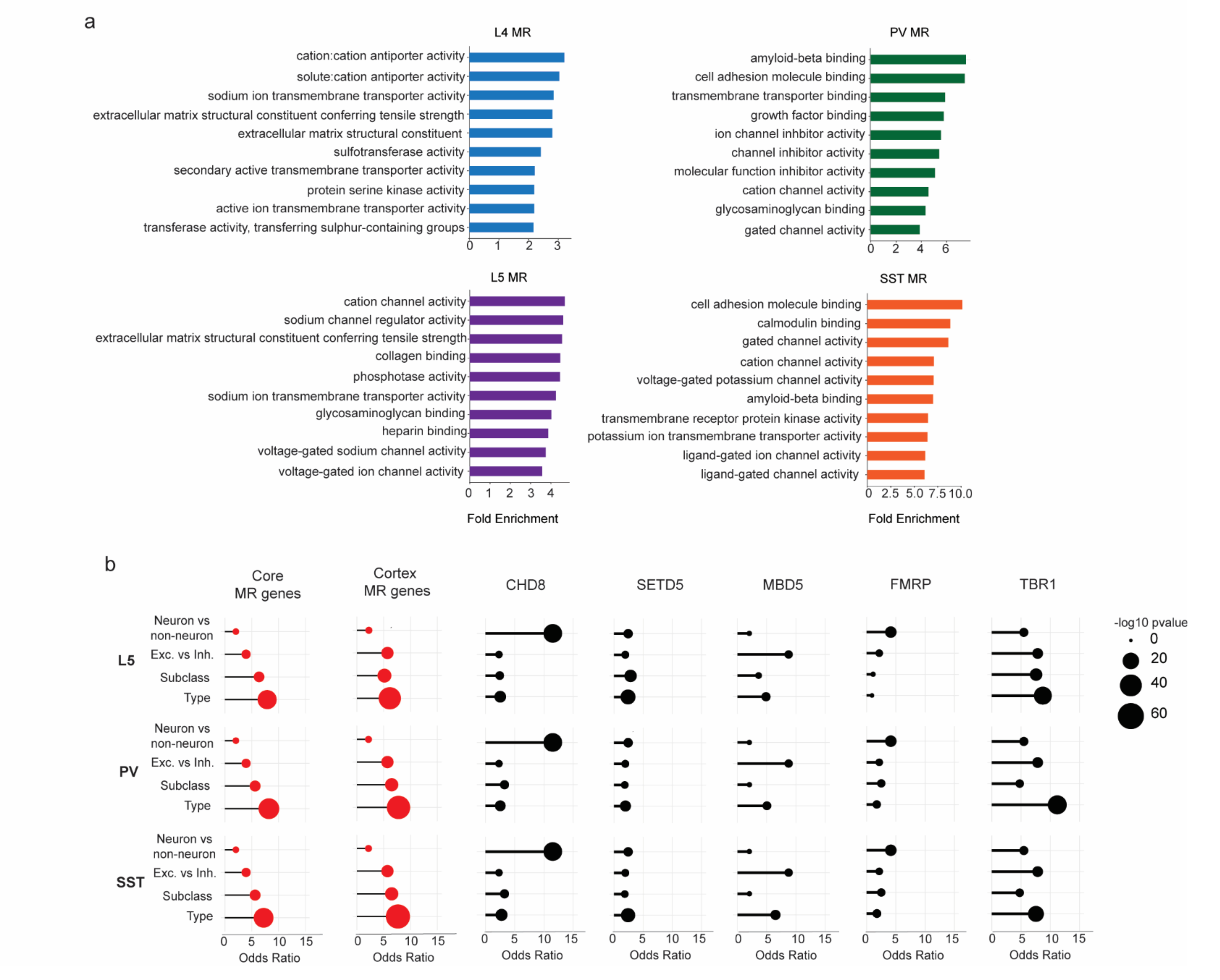
Functional annotation of subclass-defined MeCP2-repressed genes and their representation in other datasets. **a,** Gene ontology of MeCP2-repressed (MR) genes in L4, L5, PV, and SST neurons. Top 10 terms for Molecular Function shown. **b,** Overlap of differentially expressed genes across L5, PV, and SST cellular hierarchy with core MeCP2-repressed genes, MeCP2-repressed genes previously identified in the cortex^12^, and dysregulated genes detected in other NDD mouse models. Note that each differential list labeled on the left for L5, PV and SST reflects the same neuron vs. non-neuron and excitatory vs. inhibitory lists, but the lists of differential genes at the subclass and type level reflect the genes that specifically define these differentiations in L5, PV or SST cells. Analysis was performed on differentially expressed genes from INTACT RNA-seq analysis described in **Figure 1**. n=4 biological replicates per genotype per cell type, as well as the indicated published gene lists.

**Extended Data Fig. 6.**
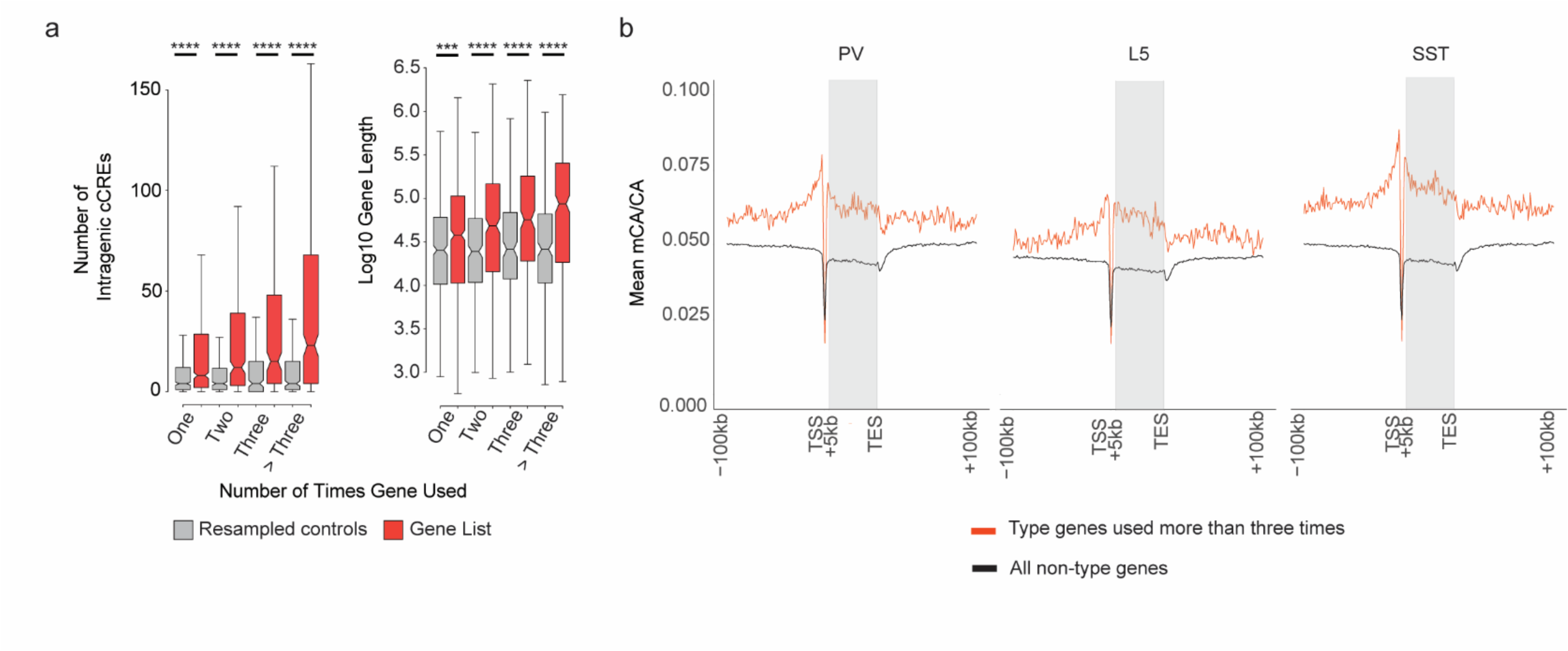
Genes that are repeatedly tuned between neuron types have characteristics that predispose them to regulation by the MeCP2 pathway. **a,** Number of intragenic cCREs (left) and gene length (right) in genes that show repeated tuning between closely related neuron types. Genes are plotted by the number of times they are detected as differential between two closely related neuron types. The center line of each boxplot is the median. Each box encloses the first and third quartile of the data. The whiskers extend to the most extreme values, excluding outliers which are outside 1.5 times the interquartile range. **b,** Aggregate profiles of mCA/CA in the cerebral cortex for genes found to be differentially expressed between closely related neuron types more than three times.

**Extended Data Fig. 7.**
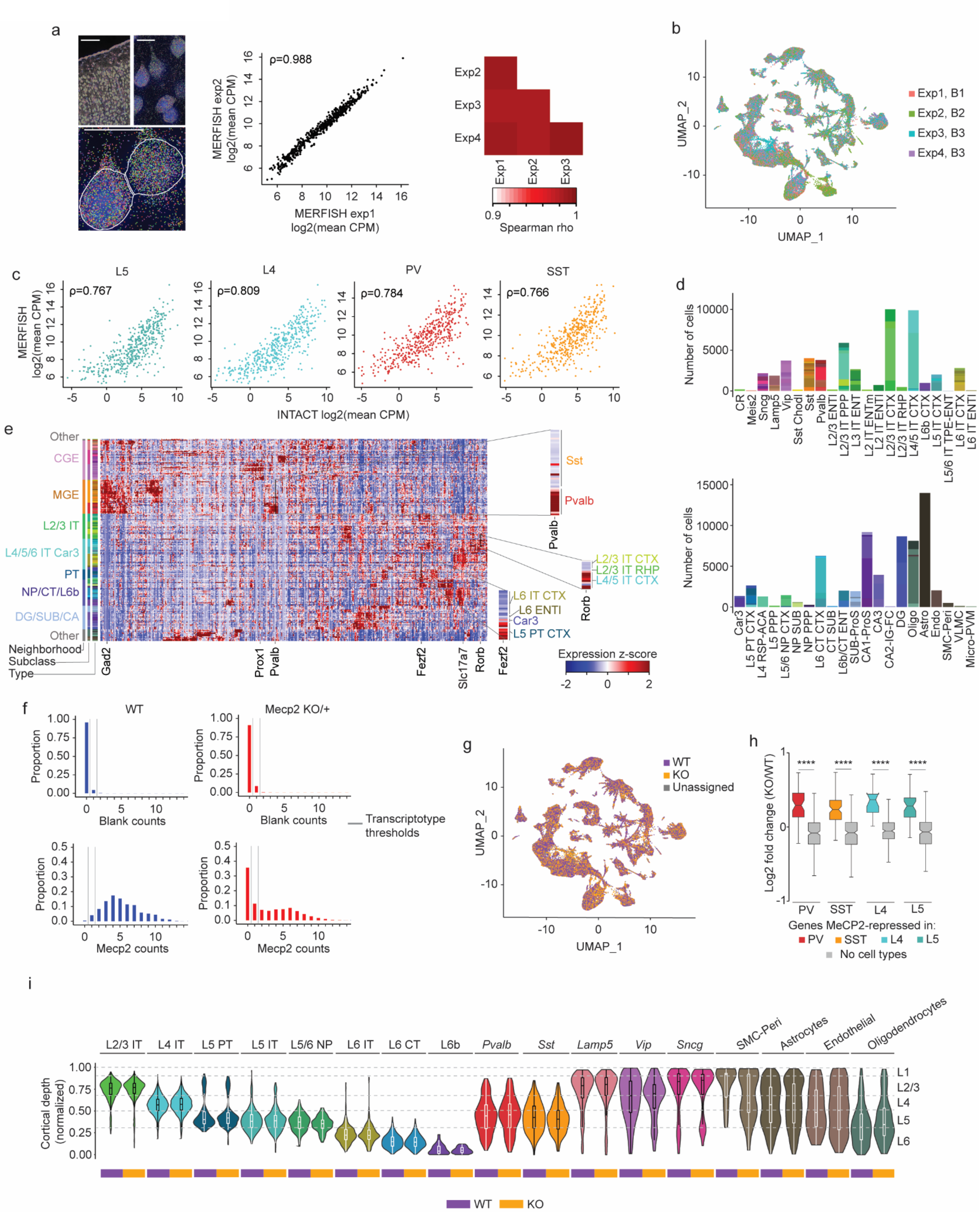
MERFISH analysis of cell types and MeCP2 expression in MeCP2 KO/+ brain shows expected gene expression and cell distributions across transcriptotypes. **a,** Left: transcripts detected (colored dots) and DAPI images of MERFISH data in one experiment. Scale bar is 250 μm for low resolution view, and 25 μm for close-ups. Middle: correlation between experiments 1 and 2 in log2 mean CPM of each gene. Right: correlation between each pair of MERFISH experiments in log2 mean CPM of each gene. ρ = Spearman’s correlation coefficient. **b,** UMAP representation of cells in MeCP2 KO/+ MERFISH experiments, colored by experiment identity and biological replicate. B1-3=biological replicate 1-3. **c,** Scatter plot of log-transformed CPMs of gene expression in neuronal subclasses (PV, SST, L4, and L5) measured by INTACT RNA-seq and MERFISH. ρ = Spearman’s correlation coefficient. **d,** Number of cells of each type in each subclass in MeCP2 KO/+ MERFISH experiments. **e.** Heatmap showing z-scores of average expression for each gene in the MERFISH panel in each cell type. Inset: close-ups of *Pvalb* (PV marker gene), *Fezf2* (L5 PT CTX marker gene), and *Rorb* (L4/5 IT CTX marker gene). **f,** Representative distributions of transcript counts per cell for *Mecp2* and a representative negative control (“Blank counts”) detected in PV cells in MERFISH analysis of wild-type or MeCP2 KO/+ coronal sections. Cut offs used for calling “WT” and “KO” transcriptotypes in the MeCP2 KO/+ are shown. **g,** UMAP representation of cells identified as either WT, KO, or unassigned in MeCP2 KO/+ MERFISH experiments. **h,** MERFISH log2 fold-change of genes identified as MeCP2-repressed in PV, SST, L4, and L5 INTACT RNA-seq analyses. ****p < 0.0001 two-sided Wilcoxon rank-sum test. The center line of each boxplot is the median. Each box encloses the first and third quartile of the data. The whiskers extend to the most extreme values, excluding outliers which are outside 1.5 times the interquartile range. **i,** Locations of distinct sublcasses of WT and KO cells across cortical layers in the visual cortex identified using MERFISH. No major differences were observed in cortical depth between WT and KO cells. The center line of each boxplot is the median. Each box encloses the first and third quartile of the data. n=3 biological replicates for MeCP2 KO/+ MERFISH across 4 imaged brain sections.

**Extended Data Fig. 8.**
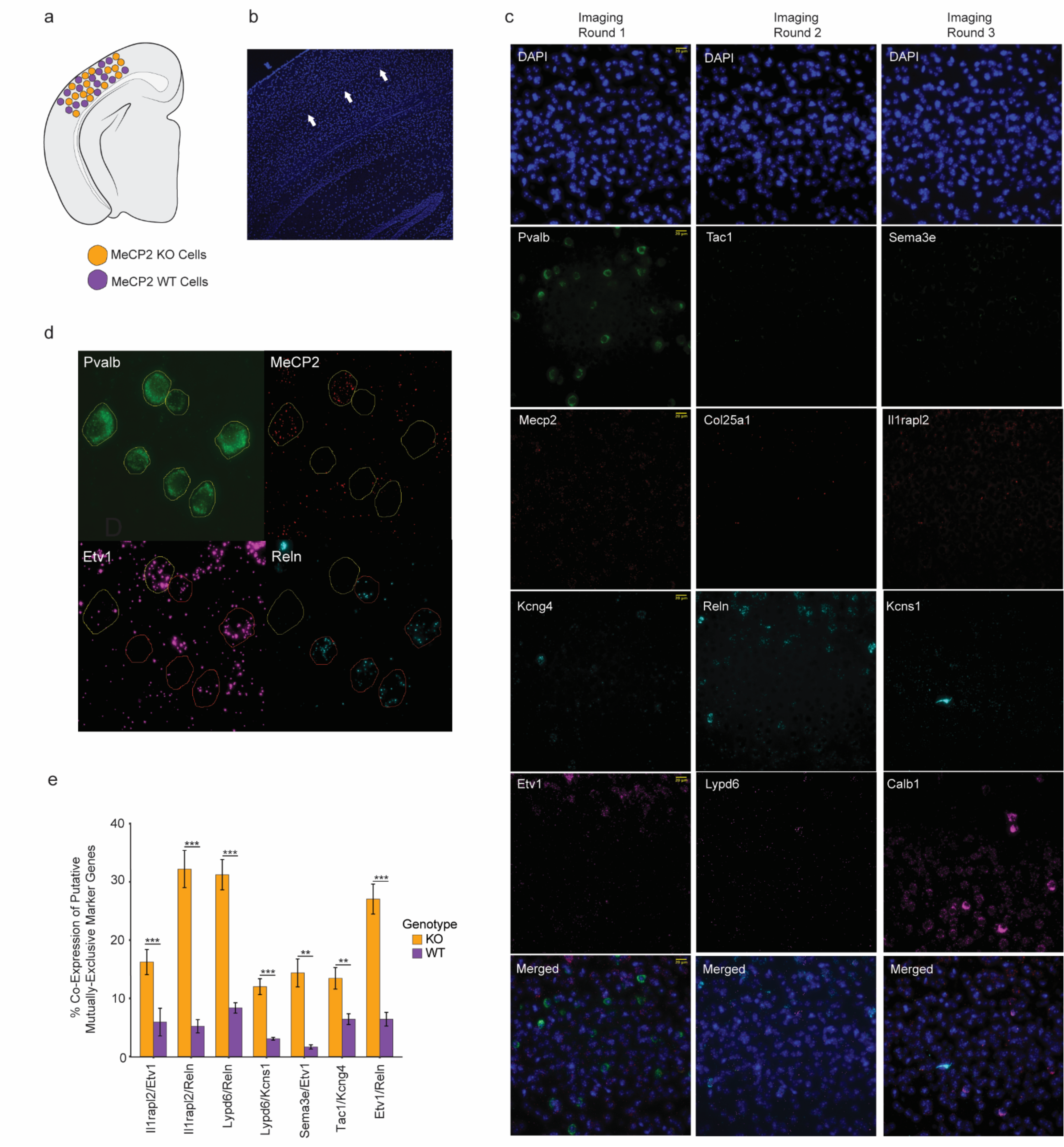
Loss of mutually exclusive type-specific gene expression in MeCP2 KO PV interneurons. **a,** MeCP2 mutant heterozygous females contain wildtype and mutant cells. **b,** Representative image of visual cortex L4 area focused on for analysis. White arrow pointing to L4 region. **c,** Representative images from all 12 target probes and DAPI stain for each of the three imaging rounds. Merged images of each round shown. **d,** RNAScope analysis of expression of mutually exclusive marker genes for visual cortex PV interneurons in MeCP2 KO and WT PV neurons. Identification of PV neurons using *Pvalb* and call of MeCP2 KO and WT cells using *Mecp2*. **e,** Bar plots of rate of co-expression of putatively mutually exclusive PV marker genes in MeCP2 null and WT PV interneurons (n=3, 50-100 cells per experiment, two-sided unpaired *t* test, **p<0.01, ***p<0.005). n=3 biological replicates for RNAScope.

**Extended Data Fig. 9.**
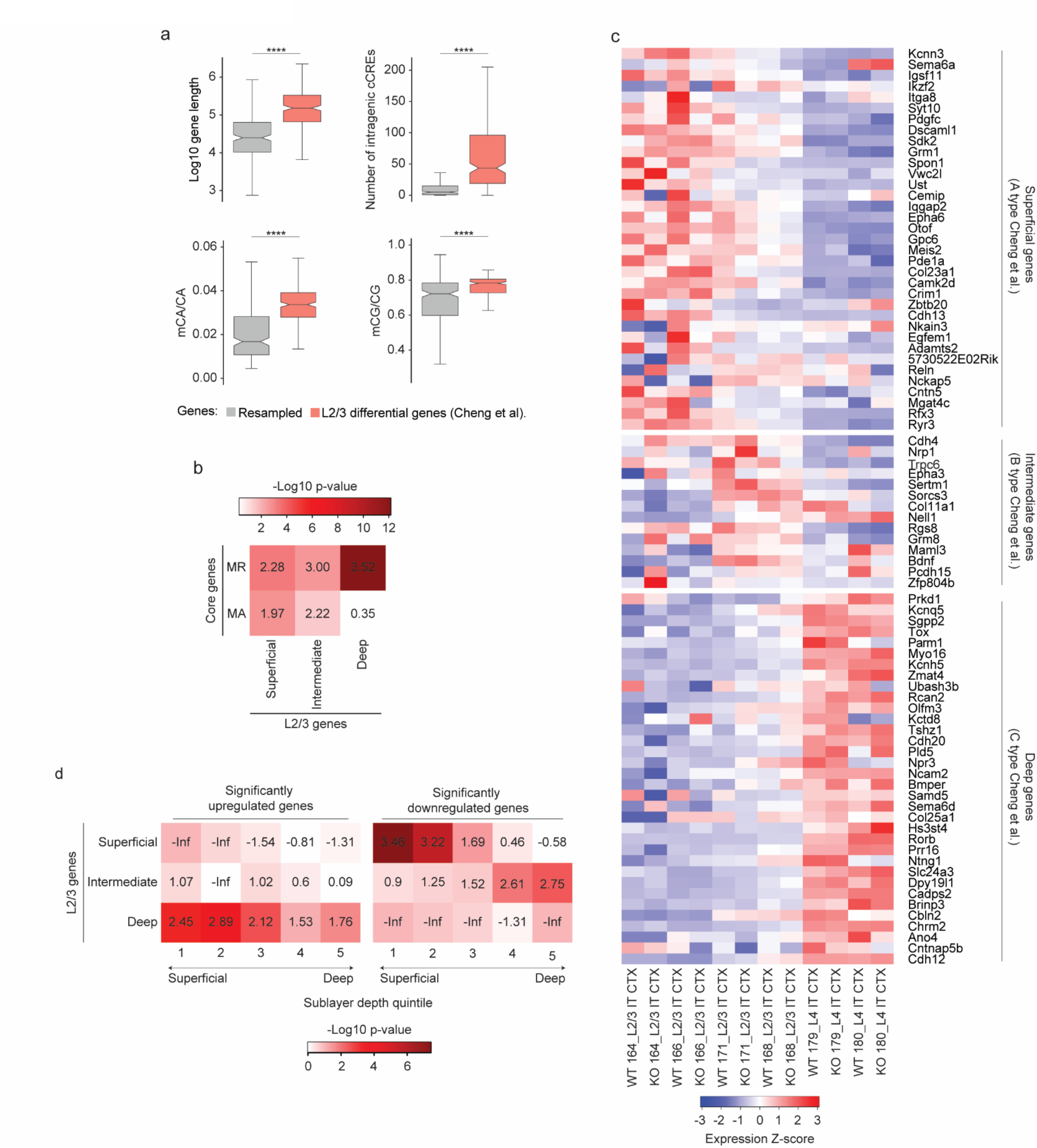
Visual cortex layer 2/3 sublayer-specific genes are targets of MeCP2 regulation. **a,** Characteristics of visual L2/3 excitatory neuron sublayer type-specific genes defined by Cheng et al., 2022^3^, showing long gene length, high numbers of intragenic cCREs and enriched mCA. ****p < 0.0001 two-sided Wilcoxon rank-sum test. **b,** Heatmap showing overlap and significance of excitatory neuron sublayer type-specific genes defined by Cheng et al., 2022^3^ with core MeCP2-regulated gene lists. Numbers in the tiles represent enrichment (log2 odds ratio) of core MeCP2-regulated genes in each L2/3 gene list. **c,** Heatmap of MERFISH-quantified expression for excitatory neuron sublayer type-specific genes defined by Cheng et al.^3^ in IT excitatory neurons in L2/3 of MeCP2 KO/+ V1 (top). **d,** Heatmap showing overlap and significance of excitatory neuron sublayer type-specific genes defined by Cheng et al.^3^ with genes detected as significantly dysregulated in pseudobulkDGE analysis of MERFISH data in excitatory neurons in sublayer depth quintiles of L2/3 of the V1. Numbers in the tiles represent enrichment (log2 odds ratio) of L2/3 genes in dysregulated gene lists of the excitatory neurons in each sublayer depth quintile of L2/3 of V1. n=3 biological replicates for MeCP2 KO/+ MERFISH across 4 imaged brain sections.

## References

1. Yao, Z. et al. A taxonomy of transcriptomic cell types across the isocortex and hippocampal formation. Cell 184, 3222–3241.e26 (2021).

2. Allaway, K. C. et al. Genetic and epigenetic coordination of cortical interneuron development. Nature 597, 693–697 (2021).

3. Cheng, S. et al. Vision-dependent specification of cell types and function in the developing cortex. Cell 185, 311–327.e24 (2022).

4. Clemens, A. W. & Gabel, H. W. Emerging Insights into the Distinctive Neuronal Methylome. Trends in Genetics 36, 816–832 (2020).

5. Lister, R. et al. Global Epigenomic Reconfiguration During Mammalian Brain Development. Science 341, 1237905 (2013).

6. de Mendoza, A. et al. The emergence of the brain non-CpG methylation system in vertebrates. Nat Ecol Evol 5, 369–378 (2021).

7. Liu, H. et al. DNA methylation atlas of the mouse brain at single-cell resolution. Nature 598, 120–128 (2021).

8. Luo, C. et al. Single-cell methylomes identify neuronal subtypes and regulatory elements in mammalian cortex. Science 357, 600–604 (2017).

9. Mo, A. et al. Epigenomic Signatures of Neuronal Diversity in the Mammalian Brain. Neuron 86, 1369–1384 (2015).

10. Boxer, L. D. et al. MeCP2 Represses the Rate of Transcriptional Initiation of Highly Methylated Long Genes. Molecular Cell 77, 294–309.e9 (2020).

11. Chen, L. et al. MeCP2 binds to non-CG methylated DNA as neurons mature, influencing transcription and the timing of onset for Rett syndrome. Proceedings of the National Academy of Sciences 112, 5509–5514 (2015).

12. Clemens, A. W. et al. MeCP2 Represses Enhancers through Chromosome Topology-Associated DNA Methylation. Mol Cell 77, 279–293.e8 (2020).

13. Gabel, H. W. et al. Disruption of DNA-methylation-dependent long gene repression in Rett syndrome. Nature 522, 89–93 (2015).

14. Guo, J. U. et al. Distribution, recognition and regulation of non-CpG methylation in the adult mammalian brain. Nat Neurosci 17, 215–222 (2014).

15. Lagger, S. et al. MeCP2 recognizes cytosine methylated tri-nucleotide and di-nucleotide sequences to tune transcription in the mammalian brain. PLOS Genetics 13, e1006793 (2017).

16. Tillotson, R. et al. Neuronal non-CG methylation is an essential target for MeCP2 function. Molecular Cell 81, 1260–1275.e12 (2021).

17. Tillotson, R. & Bird, A. The Molecular Basis of MeCP2 Function in the Brain. Journal of Molecular Biology 432, 1602–1623 (2020).

18. Stroud, H. et al. An Activity-Mediated Transition in Transcription in Early Postnatal Neurons. Neuron 107, 874–890.e8 (2020).

19. Christian, D. L. et al. DNMT3A Haploinsufficiency Results in Behavioral Deficits and Global Epigenomic Dysregulation Shared across Neurodevelopmental Disorders. Cell Reports 33, 108416 (2020).

20. Chahrour, M. et al. MeCP2, a Key Contributor to Neurological Disease, Activates and Represses Transcription. Science 320, 1224–1229 (2008).

21. Tudor, M., Akbarian, S., Chen, R. Z. & Jaenisch, R. Transcriptional profiling of a mouse model for Rett syndrome reveals subtle transcriptional changes in the brain. Proceedings of the National Academy of Sciences 99, 15536–15541 (2002).

22. Johnson, B. S. et al. Biotin tagging of MeCP2 in mice reveals contextual insights into the Rett syndrome transcriptome. Nat Med 23, 1203–1214 (2017).

23. Sugino, K. et al. Cell-Type-Specific Repression by Methyl-CpG-Binding Protein 2 Is Biased toward Long Genes. J. Neurosci. 34, 12877–12883 (2014).

24. Zhao, Y.-T., Goffin, D., Johnson, B. S. & Zhou, Z. Loss of MeCP2 function is associated with distinct gene expression changes in the striatum. Neurobiology of Disease 59, 257–266 (2013).

25. Ben-Shachar, S., Chahrour, M., Thaller, C., Shaw, C. A. & Zoghbi, H. Y. Mouse models of MeCP2 disorders share gene expression changes in the cerebellum and hypothalamus. Human Molecular Genetics 18, 2431–2442 (2009).

26. Zylka, M. J., Simon, J. M. & Philpot, B. D. Gene Length Matters in Neurons. Neuron 86, 353–355 (2015).

27. Hamagami, N. et al. NSD1 deposits histone H3 lysine 36 dimethylation to pattern non-CG DNA methylation in neurons. Molecular Cell 83, 1412–1428.e7 (2023).

28. Kurotaki, N. et al. Haploinsufficiency of NSD1 causes Sotos syndrome. Nat Genet 30, 365–366 (2002).

29. Okano, M., Bell, D. W., Haber, D. A. & Li, E. DNA Methyltransferases Dnmt3a and Dnmt3b Are Essential for De Novo Methylation and Mammalian Development. Cell 99, 247– 257 (1999).

30. Tatton-Brown, K. et al. Mutations in the DNA methyltransferase gene DNMT3A cause an overgrowth syndrome with intellectual disability. Nat Genet 46, 385–388 (2014).

31. Amir, R. E. et al. Rett syndrome is caused by mutations in X-linked MECP2, encoding methyl-CpG-binding protein 2. Nat Genet 23, 185–188 (1999).

32. del Gaudio, D. et al. Increased MECP2 gene copy number as the result of genomic duplication in neurodevelopmentally delayed males. Genetics in Medicine 8, 784–792 (2006).

33. Van Esch, H. et al. Duplication of the MECP2 Region Is a Frequent Cause of Severe Mental Retardation and Progressive Neurological Symptoms in Males. The American Journal of Human Genetics 77, 442–453 (2005).

34. Deal, R. B. & Henikoff, S. A Simple Method for Gene Expression and Chromatin Profiling of Individual Cell Types within a Tissue. Developmental Cell 18, 1030–1040 (2010).

35. Ito-Ishida, A., Ure, K., Chen, H., Swann, J. W. & Zoghbi, H. Y. Loss of MeCP2 in Parvalbumin-and Somatostatin-Expressing Neurons in Mice Leads to Distinct Rett Syndrome-like Phenotypes. Neuron 88, 651–658 (2015).

36. Liu, X. et al. Cell-Type-Specific Gene Inactivation and In Situ Restoration via Recombinase-Based Flipping of Targeted Genomic Region. J. Neurosci. 40, 7169–7186 (2020).

37. Risso, D., Ngai, J., Speed, T. P. & Dudoit, S. Normalization of RNA-seq data using factor analysis of control genes or samples. Nat Biotechnol 32, 896–902 (2014).

38. Ben-Shachar, S., Chahrour, M., Thaller, C., Shaw, C. A. & Zoghbi, H. Y. Mouse models of MeCP2 disorders share gene expression changes in the cerebellum and hypothalamus. Human Molecular Genetics 18, 2431–2442 (2009).

39. Kokura, K. et al. The Ski Protein Family Is Required for MeCP2-mediated Transcriptional Repression*. Journal of Biological Chemistry 276, 34115–34121 (2001).

40. Lyst, M. J. et al. Rett syndrome mutations abolish the interaction of MeCP2 with the NCoR/SMRT co-repressor. Nat Neurosci 16, 898–902 (2013).

41. Stroud, H. et al. Early-Life Gene Expression in Neurons Modulates Lasting Epigenetic States. Cell 171, 1151–1164.e16 (2017).

42. Li, Y. E. et al. An atlas of gene regulatory elements in adult mouse cerebrum. Nature 598, 129–136 (2021).

43. Creyghton, M. P. et al. Histone H3K27ac separates active from poised enhancers and predicts developmental state. Proceedings of the National Academy of Sciences 107, 21931– 21936 (2010).

44. Robinson, M. D., McCarthy, D. J. & Smyth, G. K. edgeR: a Bioconductor package for differential expression analysis of digital gene expression data. Bioinformatics 26, 139–140 (2010).

45. Carullo, N. V. N. & Day, J. J. Genomic Enhancers in Brain Health and Disease. Genes 10, 43 (2019).

46. Heinz, S., Romanoski, C. E., Benner, C. & Glass, C. K. The selection and function of cell type-specific enhancers. Nat Rev Mol Cell Biol 16, 144–154 (2015).

47. Heinz, S. et al. Simple Combinations of Lineage-Determining Transcription Factors Prime cis-Regulatory Elements Required for Macrophage and B Cell Identities. Molecular Cell 38, 576–589 (2010).

48. Bakken, T. E. et al. Comparative cellular analysis of motor cortex in human, marmoset and mouse. Nature 598, 111–119 (2021).

49. Gu, Y. et al. Balanced Activity between Kv3 and Nav Channels Determines Fast-Spiking in Mammalian Central Neurons. iScience 9, 120–137 (2018).

50. Miyamae, T. et al. Kcns3 deficiency disrupts Parvalbumin neuron physiology in mouse prefrontal cortex: Implications for the pathophysiology of schizophrenia. Neurobiology of Disease 155, 105382 (2021).

51. Demars, M. P. & Morishita, H. Cortical parvalbumin and somatostatin GABA neurons express distinct endogenous modulators of nicotinic acetylcholine receptors. Molecular Brain 7, 75 (2014).

52. Huntley, M. A. et al. Genome-Wide Analysis of Differential Gene Expression and Splicing in Excitatory Neurons and Interneuron Subtypes. J. Neurosci. 40, 958–973 (2020).

53. Mossink, B. et al. Cadherin-13 is a critical regulator of GABAergic modulation in human stem-cell-derived neuronal networks. Mol Psychiatry 27, 1–18 (2022).

54. Paul, A. et al. Transcriptional Architecture of Synaptic Communication Delineates GABAergic Neuron Identity. Cell 171, 522–539.e20 (2017).

55. Tasic, B. et al. Shared and distinct transcriptomic cell types across neocortical areas. Nature 563, 72–78 (2018).

56. Chen, K. H., Boettiger, A. N., Moffitt, J. R., Wang, S. & Zhuang, X. Spatially resolved, highly multiplexed RNA profiling in single cells. Science 348, aaa6090 (2015).

57. Kim, E. J. et al. Extraction of Distinct Neuronal Cell Types from within a Genetically Continuous Population. Neuron 107, 274–282.e6 (2020).

58. Zhang, M. et al. Spatially resolved cell atlas of the mouse primary motor cortex by MERFISH. Nature 598, 137–143 (2021).

59. Zhang, Z. et al. Epigenomic diversity of cortical projection neurons in the mouse brain. Nature 598, 167–173 (2021).

60. Renthal, W. et al. Characterization of human mosaic Rett syndrome brain tissue by single-nucleus RNA sequencing. Nat Neurosci 21, 1670–1679 (2018).

61. Hao, Y. et al. Integrated analysis of multimodal single-cell data. Cell 184, 3573–3587.e29 (2021).

62. Lun, A. T. L., McCarthy, D. J. & Marioni, J. C. A step-by-step workflow for low-level analysis of single-cell RNA-seq data with Bioconductor. Preprint at 10.12688/f1000research.9501.2 (2016).

63. Kishi, N. & Macklis, J. D. MECP2 is progressively expressed in post-migratory neurons and is involved in neuronal maturation rather than cell fate decisions. Mol Cell Neurosci 27, 306–321 (2004).

64. Kishi, N. & Macklis, J. D. MeCP2 functions largely cell-autonomously, but also non-cell-autonomously, in neuronal maturation and dendritic arborization of cortical pyramidal neurons. Exp Neurol 222, 51–58 (2010).

65. Chao, H.-T. et al. Dysfunction in GABA signalling mediates autism-like stereotypies and Rett syndrome phenotypes. Nature 468, 263–269 (2010).

66. Goertsen, D. et al. AAV capsid variants with brain-wide transgene expression and decreased liver targeting after intravenous delivery in mouse and marmoset. Nat Neurosci 25, 106–115 (2022).

67. Graybuck, L. T. et al. Enhancer viruses for combinatorial cell-subclass-specific labeling. Neuron 109, 1449–1464.e13 (2021).

68. Vormstein-Schneider, D. et al. Viral manipulation of functionally distinct interneurons in mice, non-human primates and humans. Nat Neurosci 23, 1629–1636 (2020).

69. Tasic, B. et al. Adult mouse cortical cell taxonomy revealed by single cell transcriptomics. Nat Neurosci 19, 335–346 (2016).

70. Wu, S. J. et al. Cortical somatostatin interneuron subtypes form cell-type-specific circuits. Neuron 111, 2675–2692.e9 (2023).

71. Cusanovich, D. A. et al. A Single-Cell Atlas of In Vivo Mammalian Chromatin Accessibility. Cell 174, 1309–1324.e18 (2018).

72. Nord, A. S. & West, A. E. Neurobiological functions of transcriptional enhancers. Nat Neurosci 23, 5–14 (2020).

73. Visel, A. et al. A High-Resolution Enhancer Atlas of the Developing Telencephalon. Cell 152, 895–908 (2013).

74. Dehorter, N. et al. Tuning of fast-spiking interneuron properties by an activity-dependent transcriptional switch. Science 349, 1216–1220 (2015).

75. Di Bella, D. J. et al. Molecular logic of cellular diversification in the mouse cerebral cortex. Nature 595, 554–559 (2021).

76. Mayer, C. et al. Developmental diversification of cortical inhibitory interneurons. Nature 555, 457–462 (2018).

77. Mardinly, A. R. et al. Sensory experience regulates cortical inhibition by inducing IGF1 in VIP neurons. Nature 531, 371–375 (2016).

78. Cohen, S. et al. Genome-Wide Activity-Dependent MeCP2 Phosphorylation Regulates Nervous System Development and Function. Neuron 72, 72–85 (2011).

79. Dobin, A. et al. STAR: ultrafast universal RNA-seq aligner. Bioinformatics 29, 15–21 (2013).

80. Quinlan, A. R. & Hall, I. M. BEDTools: a flexible suite of utilities for comparing genomic features. Bioinformatics 26, 841–842 (2010).

81. Love, M. I., Huber, W. & Anders, S. Moderated estimation of fold change and dispersion for RNA-seq data with DESeq2. Genome Biology 15, 550 (2014).

82. Rao, S. S. P. et al. A 3D Map of the Human Genome at Kilobase Resolution Reveals Principles of Chromatin Looping. Cell 159, 1665–1680 (2014).

83. Bonev, B. et al. Multiscale 3D Genome Rewiring during Mouse Neural Development. Cell 171, 557–572.e24 (2017).

84. Moffitt, J. R. et al. High-throughput single-cell gene-expression profiling with multiplexed error-robust fluorescence in situ hybridization. Proceedings of the National Academy of Sciences 113, 11046–11051 (2016).

85. Hafemeister, C. & Satija, R. Normalization and variance stabilization of single-cell RNA-seq data using regularized negative binomial regression. Genome Biol 20, 296 (2019).

86. McInnes, L., Healy, J., Saul, N. & Großberger, L. UMAP: Uniform Manifold Approximation and Projection. Journal of Open Source Software 3, 861 (2018).

87. Morimoto, M. M., Uchishiba, E. & Saleem, A. B. Organization of feedback projections to mouse primary visual cortex. iScience 24, 102450 (2021).

88. Shamash, P., Carandini, M., Harris, K. & Steinmetz, N. A tool for analyzing electrode tracks from slice histology. 447995 Preprint at 10.1101/447995 (2018).

89. Wang, Q. et al. The Allen Mouse Brain Common Coordinate Framework: A 3D Reference Atlas. Cell 181, 936–953.e20 (2020).

